# GSK3β-SCF^FBXW7^ mediated phosphorylation and ubiquitination of IRF1 are required for its transcription-dependent turnover

**DOI:** 10.1101/481911

**Authors:** Alexander J. Garvin, Ahmed H.A. Khalaf, Alessandro Rettino, Jerome Xicluna, Laura Butler, Joanna R. Morris, David M. Heery, Nicole M. Clarke

## Abstract

IRF1 (Interferon Regulatory Factor-1) is the prototype of the IRF family of DNA binding transcription factors. IRF1 protein expression is regulated by transient up-regulation in response to external stimuli followed by rapid degradation via the ubiquitin-proteasome system. Here we report that DNA bound IRF1 turnover is promoted by GSK3β (Glycogen Synthase Kinase 3β) via phosphorylation of the T181 residue which generates a phosphodegron for the SCF (Skp-Cul-Fbox) ubiquitin E3-ligase receptor protein Fbxw7α (F-box/WD40 7). This regulated turnover is essential for IRF1 activity, as mutation of T181 results in an improperly stabilised protein that accumulates at target promoters but fails to induce RNA-Pol-II elongation and subsequent transcription of target genes. Consequently, the anti-proliferative activity of IRF1 is lost in cell lines expressing T181A mutant. Further, cell lines with dysfunctional Fbxw7 are less sensitive to IRF1 overexpression, suggesting an important co-activator function for this ligase complex. As T181 phosphorylation requires both DNA binding and RNA-Pol-II elongation, we propose that this event acts to clear “spent” molecules of IRF1 from transcriptionally engaged target promoters.

## INTRODUCTION

IRF1 is a transcription factor essential for regulating a number of cellular responses including, immunity, apoptosis and DNA repair (1-4). IRF1 is highly modified by several post-translational modifications. Phosphorylation of a cluster of residues in the C terminus by casein kinase II may be required for activity as mutation of these residues reduces reporter activity (5). These residues overlap with sites reported to be targeted by IKKε, although mutation of these sites has little impact on IRF1 luciferase reporter activity they are needed for synergism with RelA (6). IRF1 is also phosphorylated on Y109 in the DBD (DNA binding domain). This modification plays a role in dimerisation with IRF8 and transcriptional activity (7). IRF1 also undergoes a number of other modifications, including SUMOylation (8) methylation (9) and acetylation (10).

IRF1 is a highly unstable protein with a half-life of around 30 minutes (11) that can be stabilised through interaction with the chaperone Hsp90 (12). Several reports have studied the ubiquitin-dependent regulation of IRF1 turnover (13-15), with roles for MDM2 and CHIP (C-terminus of HSC70 interacting protein) E3 ligases in ubiquitination of IRF1 reported. In these studies, IRF1 is modified by ubiquitin polymers formed through both K48 and K63 linkages (13-17). While a role for ubiquitination in the proteasome-mediated degradation of IRF1 is clear, little is known regarding what signals ubiquitination of IRF1 and if turnover regulates IRF1 transcriptional activity beyond regulating abundance.

Crosstalk between phosphorylation and the ubiquitin machinery is important for regulating protein abundance, activity and interactions (18,19). In some contexts phosphorylation generates PTM motifs (phospho-degrons) that are recognised by receptor proteins associated with the ubiquitin-proteasome degradation machinery. The activities of multiple transcription factors are regulated by this type of cross-talk (19). Consequently phosphorylation can serve as an important regulatory switch in target ubiquitination and degradation. GSK3β is a serine/threonine kinase with a preference for a +4 “priming” phosphorylated or acidic substrate residue for effective catalysis. Many transcription factors targeted for phosphorylation-mediated degradation are GSK3β substrates, in concert with the Fbxw7, a SCF (Skp-Cul-Fbox) phospho-substrate receptor protein (20-24). GSK3β is known to play a role in cancer and has been documented as having both, cancer promoting and cancer inhibiting functions. Together with GSK3β, Fbxw7 controls the turnover of a number of key oncogenes such as c-myc, Cyclin E and NOTCH (25-29). and has emerged as an important tumour suppressor that is frequently mutated in cancer (30).

While extensively modified, we understand relatively little about how IRF1 activity is modulated at the post translational level. In this study we focused on a pair of previously uncharacterised phosphorylation sites and uncovered a novel mechanism by which cells mark IRF1 as “spent” at the end of the transcriptional cycle.

## MATERIAL AND METHODS

### Cell lines, siRNA, antibodies and chemicals

Cells were maintained in the recommended growth media supplemented with 10% FBS, 50U/mL Penicillin-Streptomycin and 2mM L-glutamine (Supplementary Table 1). H3396 doxycycline-inducible stable cell lines were generated using pCDNA6-TetR system (Invitrogen) and pCDNA4-murine IRF1 or vector alone and selected with Zeocin. Doxycycline was used at 2µg/mL for indicated time points. Dharmacon siRNA pools were used throughout the study. All siRNA were used at 10nM final concentration for knockdown. Transfection of siRNA was performed with InterFerin (Polyplus). MG132, DRB, Doxycyline and CHX were from Sigma Aldrich, GSK3 inhibitors BIO and methyl-BIO were from Merck. Details of antibodies used can be found in supplementary table 2. The details of primers used can be found in supplementary table 3.

### Luciferase reporter assay, Cycloheximide chase assay

Reporter assays; cells were seeded for 24 hours in 24 well plates followed by transfection with reporter construct, IRF1 and internal control CMV-βGAL. Lysis was carried out 48 hours post-transfection essentially according to manufacturer’s instructions (Applied Biosystems). Luminescence was detected on a Berthold Orion micro-plate luminometer. For analysis of protein degradation, cycloheximide (CHX Sigma Aldrich) chase assays were performed as follows; cells were seeded on 6 well plates for 24 hours, transfected with 2.5µg/well of IRF1 and 24 hours later cells treated with 25 µg/mL CHX for the indicated times followed by lysis and immunoblot against IRF1 and β-actin loading control.

### Immunoprecipitations, Ubiquitination assays, GST-pulldown assays

For immunoprecipitations, cell lysates (0.5mg) were diluted in RIPA buffer (50mM Tris-HCl, pH 7.5, 150Mm NaCl, 1% NP40, 0.1% SDS, 0.5% sodium deoxycholate, 1mM EDTA) supplemented with protease and phosphatase inhibitor cocktail (SIGMA), 1mM DTT and 100mM N-Ethylmaleimide (NEM). After pre-clearing for 1 hour with Protein G beads, the lysates were incubated overnight at 4°C with the appropriate antibody, washed three times with RIPA buffer and eluted in loading buffer with boiling. Co-immunoprecipitations (1mg lysate) between FLAG-IRF1 and GSK3β-HA were carried out as above but with TNE buffer (10mM Tris (pH 7.4), 1mM EDTA and 200mM NaCl). For GST-F-box co-immunoprecipitations extracts were made in NP40 lysis buffer (50mM Tris pH 7.5, 150mM NaCl, 0.5% NP40 and phosphatase/protease inhibitors). 0.5mg of lysate was incubated at 4°C with glutathione – sepharose beads (GE Healthcare) for 3 hours, followed by 3x washes in NP40 buffer and elution in loading buffer.

For *in vitro* pulldown assays, GST or GST-IRF1 (1µg) conjugated GSH beads were diluted in NETN buffer (20mM Tris-HCl pH 8, 100mM NaCl, 1mM EDTA, 0.5% NP40 and phosphatase and protease inhibitor cocktail). In vitro translated 35S-labelled proteins (TnT Promega) were added to GST beads the beads overnight at 4°C followed by 3x washes with NETN. Boiled eluates were separated by SDS-PAGE and the gel was fixed in fixing solution (10% acetic acid 10% methanol) for 30 minutes before being treated with amplifier solution (Amersham) for 30 minutes with gentle rocking. Gels were dried and exposed to film at −80°C.

For ubiquitination assays, 60% confluent 10cm dishes of HEK293 were transfected with 6xHis-myc-Ubiquitin (2.5 µg) and FLAG IRF1 (2.5µg). 40 hours later MG132 (10 µM) was added for 5 hours. Duplicate transfected plates were treated with 0.01% DMSO as vehicle control. 6xHis nickel pulldowns were carried out essentially as described (31). Lysate fractionation was performed as described (32).

### In vitro-kinase assays

Recombinant GSK3β (New England Biolabs) was diluted to 20ng/µL in kinase buffer and incubated with 4µg of GST-IRF1. The final concentration of ATP was 250 µM, in 25 µL. The reaction was carried out at 37°C for 1 hour, and terminated by the addition of 6µL of 5X Laemmli buffer (with 100mM DTT).

### RNA extraction, reverse transcription, real-time PCR and ChIP

All RNA extractions, reverse transcription, real-time PCR reactions and chromatin immunoprecipitations were performed essentially as in [2]. All primer sequences are available upon request.

### Proliferation assays

Proliferation assays were performed in H3396 stable cell lines expressing murine IRF1 in pBabeSIN puro under the control of a tetracycline inducible promoter. These vectors were used to transduce H3396 together with a retrovirus encoding the Tet transactivator in a Tet-on configuration selectable with hygromycin and a Tet-repressor, which consists of a Tet DNA binding domain and a KRAB domain selectable with neomycin in order to minimise the background expression. Proliferation assays were carried out by plating the stables at 300 cells per well on a 96-well plate and then treating with doxycycline at 3 µg/mL for the indicated times. Cells were then trypsinized, resuspended in PBS/trypan blue and counted by light microscope using the trypan blue exclusion method. All cell counts were done in quadruplicate. For proliferation of other lines, cells were transduced with pBabeSIN puro, IRF1-WT or IRF1 T181A for 48 hours prior to selection with 1ug/mL puromycin to remove non-transduced cells. To assess clonal growth, cells were counted and plated at low dilution (100-500 cells) on 48 well plates and left to grow for 10 days prior to staining with crystal violet 0.05% in 50% methanol). For short-term growth assays, cells were plated in 24 well plates following puromycin selection and allowed to grow for 4 days. Cells were counted in triplicate after trypan blue staining.

### Statistics

Unless stated otherwise all quantitative assays (reporter, QPCR and ChIP) were performed three times with three technical repeats.* denotes statistical significance with a *p* value less than 0.05 as detected by Students t-test ** <0.01, *** <0.005.

## RESULTS

### IRF1 is phosphorylated by GSK3β

In a search for potential IRF1 phosphorylation sites we focused on a sequence within the IRF1 transactivation domain (TAD) that shares similarity to a subset of consensus GSK3 target sites. This sequence is conserved in mammalian species and comprises a threonine residue (T181 in human IRF1) with an upstream “priming” residue (S185) (Fig.1A, S1A/B). To determine if IRF1 could be directly phosphorylated by GSK3β we performed *in vitro* kinase assays with recombinant GSK3β and murine GST-IRF1. The reaction mix was subjected to western blotting using an antibody raised against phospho-threonine-proline dipeptide. As murine IRF1 contains only a single threonine-proline dipeptide sequence, we could therefore use this antibody to assess IRF1 T181 phosphorylation. In addition to a migration shift in the GST-IRF1 protein, we detected phosphorylation of IRF1 after exposure to GSK3β indicating that GSK3β could directly modify IRF1 (Fig.1B). To support this result, we next transfected HEK293 cells (which lack detectable endogenous IRF1 protein) with expression vectors for FLAG-IRF1 (mouse) and GSK3β-HA. Lysates were immunoprecipitated with the anti-phospho-TP antibody and blotted with anti-FLAG antibody. In cells transfected with FLAG-IRF1 only, we were able to detect phosphorylation of IRF1 suggesting that endogenous GSK3 kinases or other kinases can target this residue. However this signal was strongly increased upon over-expression of WT GSK3β-HA providing further evidence that IRF1 T181 could be targeted by GSK3β (Fig. 1C).

**Figure 1.**
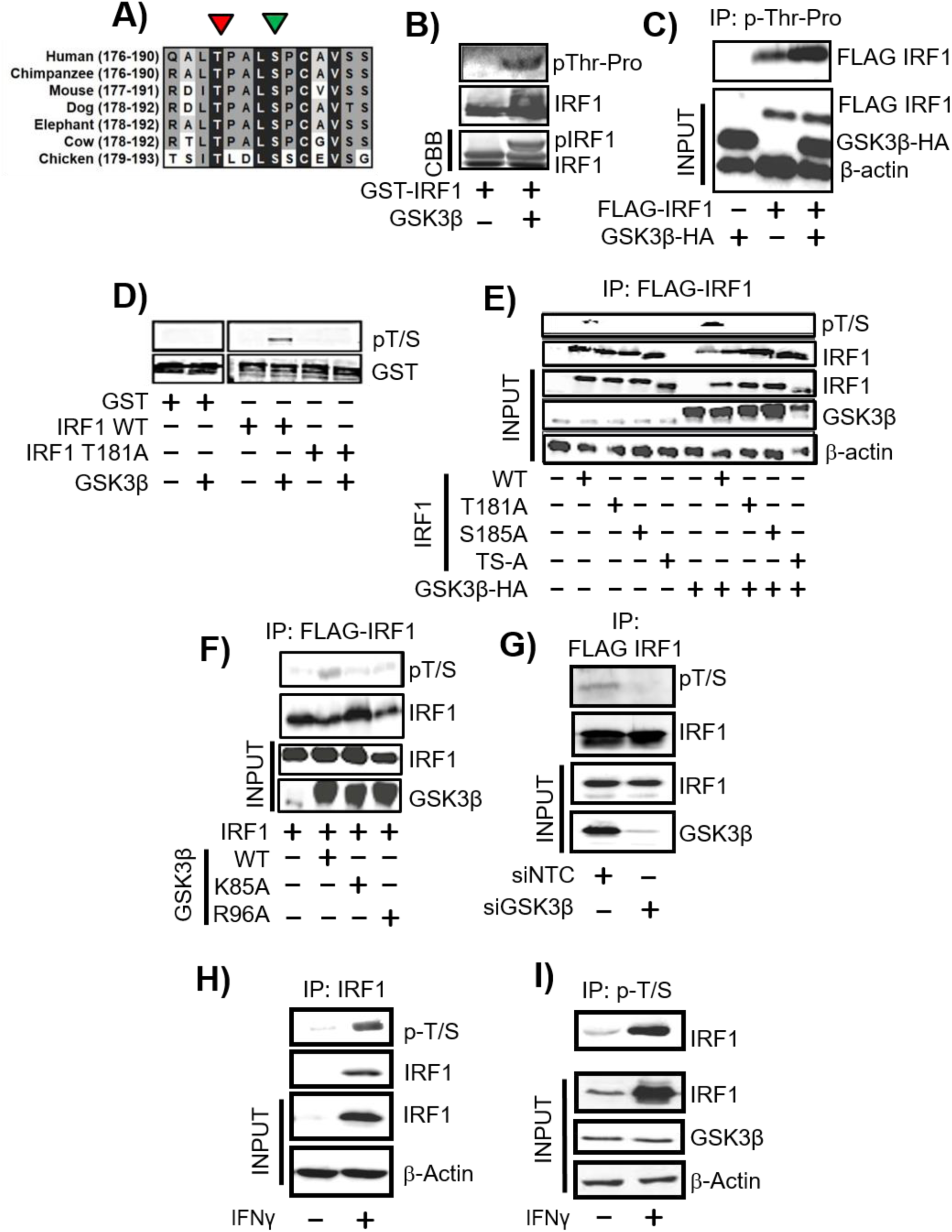
IRF1 is phosphorylated by GSK3β. **A**) Sequence conservation of the putative GSK3β phospho-target sequence in different species. The phosphorylated T180 and the +4 priming site (S184) (T181/S185 in murine sequence) residues are indicated by red and green arrows, respectively. Residue numbers in parenthesis. **B**) *In vitro* kinase assay performed using recombinant GSK3β and purified GST-IRF1 protein as substrate. The reaction products were resolved by SDS PAGE and GST-IRF1 T181 phosphorylation revealed by western blotting using anti-pTP antibody (top panel). Altered migration of GST-IRF1 after phosphorylation by GSK3β was also visible after western blot detection using anti-IRF1 (middle panel), or by Coomassie brilliant blue (CBB) staining (bottom panel). **C**) Lysates from HEK293 expressing GSK3β-HA and mouse FLAG-IRF1, immunoprecipitated with the p-TP antibody. Immunoprecipitated IRF1 was detected with anti-FLAG antibody. Inputs (10%) indicate the expression of transfected FLAG-IRF1 and GSK3β-HA proteins, and loading control β-actin. **D**) *In vitro* kinase assay performed as in 1B but visualised by immunoblot with pT/S (pThr^58^/Ser^62^ c-myc) and with GST to show loading. The T181A mutant is included to demonstrate specificity of the antibody. **E**) of HEK293 expressing FLAG-IRF1 WT or mutants together with GSK3β-HA or empty vector were immunoprecipitated with anti-FLAG beads. IRF1 T181/S185 dual phosphorylation was detected by western blotting with pT/S antibody (top panel). Successful IP of IRF1 proteins in the extracts was confirmed by re-probing with anti-FLAG antibody (second panel). Inputs (10%) are shown in the lower three panels and indicate the levels of IRF1 (anti-FLAG), GSK3β (anti-HA) and loading control β-actin. **F**) As for F, but with GSK3β kinase inactive (K85A) and priming mutants (R96A). **G**) HEK293 were siRNA depleted for control of GSK3β for 24 hours prior to transfection with FLAG-IRF1 for a further 48 hours. Lysates were immunoprecipitated and probed with pT/S antibody and FLAG to show IP efficiency. **H**) Extracts from H3396 cells treated for 3 hours with IFNγ (1000U/mL) or vehicle were immunoprecipitated with anti-pT/S and probed with IRF1. Input lysates (10%) are shown below against IRF1, GSK3β and actin **I**) MRC5 lysates immunoprecipitated with IRF1 followed by probe with p-T/S. Note absence of detectable IRF1 protein in un-treated cells.

We noted that the sequence surrounding T181 (i.e. 181-TPALSP-186) is conserved in a subset of phosphoproteins (Fig. S1A) including the transcription factor c-Myc (58-TPPLSP-63) and for which there are commercially available phospho-antibodies. Indeed, we were able to demonstrate that an antibody raised against phospho-T58/S62 (pT/S) of c-MYC also detected wild type FLAG-IRF1 in a cold kinase assay but not a FLAG-IRF1 T181A mutant (Fig. 1D). To further confirm that this antibody detects IRF1 phosphorylated at T181, we co-transfected HEK293 cells with GSK3β-HA or empty vector in combination with Flag-IRF1 WT, T181A, S185A or T181A/S185A vectors (Fig. 1E). Following immunoprecipitation, FLAG-IRF1 proteins were blotted with the pT/S antibody. Again, while an increase in phosphorylation of wild type IRF1 was detected on GSK3β over-expression, this antibody failed to detect IRF1 proteins containing T181A, S185A or both in combination. Similar results were obtained using a different tag for detection i.e. YFP-IRF1 (Fig. S1C). Over-expression of catalytic mutant GSK3β (K85A) did not promote increased phosphorylation of IRF1, neither did the R96A “priming” mutant of GSK3β that cannot phosphorylate residues if a +4 priming site is already phosphorylated, suggesting that S185 is a likely to function as a priming residue (Fig. 1F). Further the double alanine mutant of IRF1 migrates more rapidly suggesting modification of both residues. Additionally we confirmed GSK3β was the dominant kinase for this site using siRNA depletion followed by detection of phosphorylation of immunoprecipitated FLAG-IRF1 with the pT/S antibody (Fig. 1G)

### Endogenous IRF1 protein is phosphorylated by GSK3β

We next determined that pT/S antibody could immunoprecipitate endogenous human IRF1 in H3396 breast cancer cells in which IRF1 expression was increased by IFNγ treatment. (Fig. 1H). Similar experiments were performed in MRC5 fibroblasts in which the pT/S antibody successfully detected the immunoprecipitated IRF1 (Fig.1I).

Taken together, these data provide strong evidence that both exogenous murine and endogenous human IRF1 proteins are subjected to dual Thr^180^/Ser^184^ phosphorylation by GSK3β.

### IRF1 interacts with GSK3β

To explore whether IRF1 interacts with GSK3β, co-immunoprecipitations were performed on extracts of HEK293 cells overexpressing FLAG-IRF1 and GSK3β-HA proteins. We successfully co-precipitated IRF1/GSK3β complexes using either epitope tag (Fig. 2A/B). Importantly we were also able to detect reciprocal co-IP of endogenous IRF1 and GSK3β present in H3396 cells (Fig. 2C). These complexes were only detectable after treatment of H3396 cells with the proteasome inhibitor MG132, suggesting that the interaction may be associated with degradation of IRF1. Interaction of these proteins *in vitro* was confirmed by GST-IRF1 pulldown of ^35^[S}-methionine labelled GSK3β (Fig. 2D). Thus, the interaction between GSK3β and IRF1 supports our hypothesis that IRF1 is a phosphorylation target of GSK kinases.

**Figure 2.**
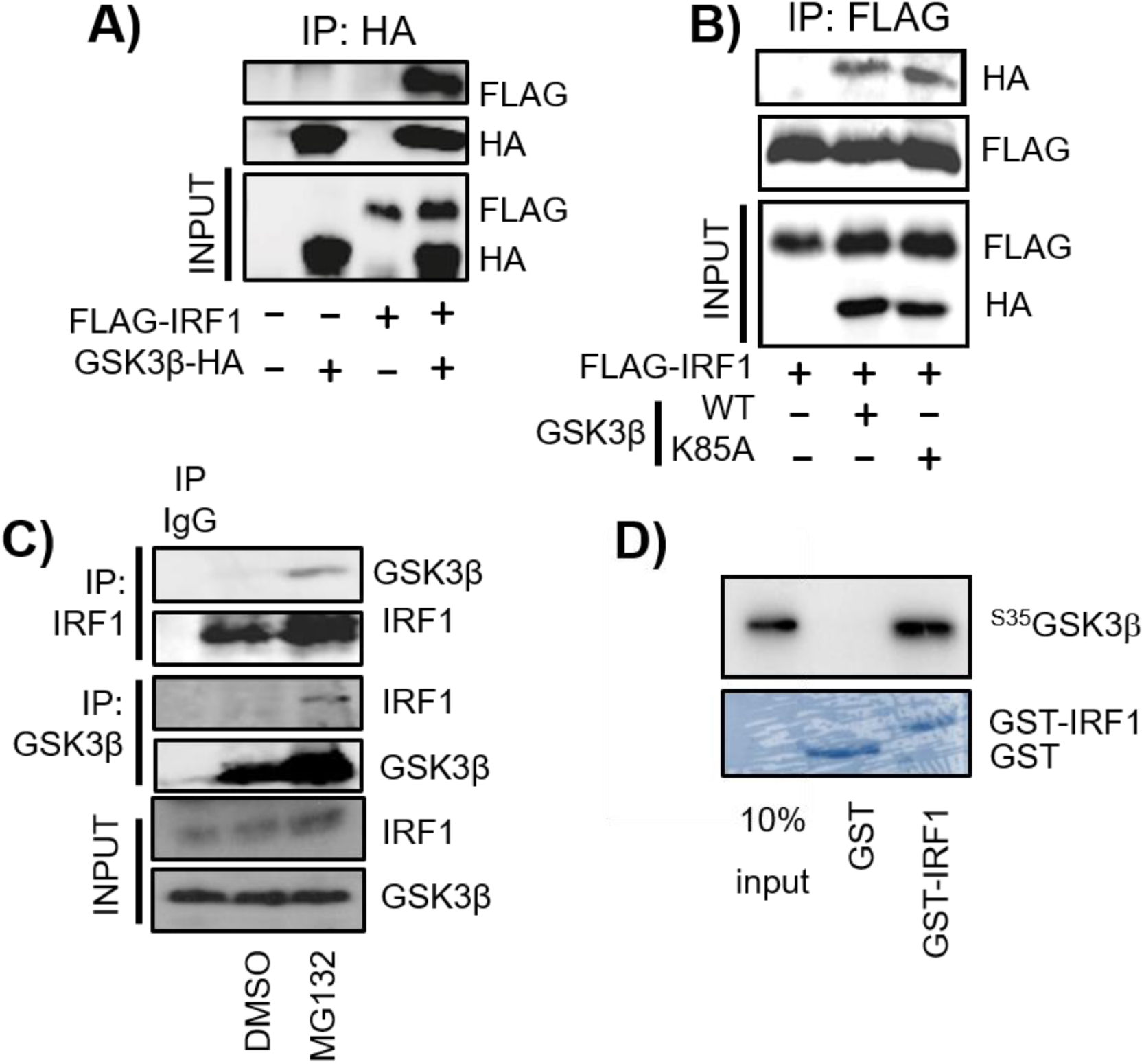
IRF1 interacts with GSK3β. **A**) Extracts from HEK293 cells expressing FLAG-IRF1 and GSK3β-HA were immunoprecipitated with anti-HA. Shown are western blots used to reveal co-precipitated HA-GSK3β or FLAG-IRF1 proteins. Expression levels in the inputs (10%) are shown in the bottom panels. **B**) As for A, but using anti-FLAG immunoprecipitation and including kinase inactive GSK3β-HA (K85A) **C**) Extracts from H3396 cells pre-treated with MG132 (10µM/ 3 hr) or DMSO, immunoprecipitated with IgG, anti-IRF1 or anti-GSK3β antibodies to reveal endogenous complexes. **D**) GST pulldown experiment using bacterially expressed, partially purified GST-IRF1 immobilised on glutathione sepharose beads and incubated with *in vitro* transcribed/translated ^35^[S]-methionine-labelled GSK3β-HA. Proteins retained on the beads were visualized by autoradiography. 10% input of radiolabeled product is shown as input.

### GSK3β is required for IRF1 transcriptional activity

To investigate the effects of phosphorylation on IRF1 activity, we measured IRF1-dependent transactivation in reporter assays. For overexpression we used Cos7 cells that lack detectable IRF1, allowing us to discount possible effects of endogenous IRF1 proteins. Our results showed that IRF1 reporter activity on the TRAIL / TNFSF10 (TNFα Related Apoptosis Inducing Ligand) promoter fragment was reduced following treatment with GSK3 inhibitors LiCl or Inhibitor-BIO but not the inactive derivative methyl-BIO (Fig. 3A&B). This suggested that GSK3 is required for IRF1-dependent stimulation of reporter activity. Next, we overexpressed GSK3β in TRAIL promoter reporter assays, but found no increase in IRF1 activity when GSK3β was over-expressed, however we did detect a dose-dependent decrease in IRF1 activity when the K85A (kinase inactive) mutant was overexpressed (Fig. 3C). Given that this mutant interacts with IRF1, but does not phosphorylate it, it is likely to be a dominant negative effect. GSK3 inhibitors do not discriminate between the two GSK3 isoforms (α/β) and can also inhibit other related kinases. To verify the requirement of GSK3β in IRF1 transcriptional activity we used siRNA directed against GSK3β. Reporter assays revealed that knockdown of GSK3β significantly reduced IRF1-mediated activation of the TRAIL promoter (Fig. 3D). These results demonstrate that GSK3β is required for the full transcriptional function of IRF1, but over-expressing WT GSK3β does not potentiate IRF1 activity further. We then wanted to determine if phosphorylation of residues (Thr^181^/Ser^185^) are required for IRF1 activity. Expression vectors for wild type or mutant IRF1 proteins were co-transfected with the TRAIL reporter or 4xISRE (Interferon Stimulated Response Element) reporter constructs in Cos7 cells. All three of the phosphorylation-deficient mutants displayed reduced activity in the reporter assays (Fig. 3E) despite being expressed at similar levels compared to wild type and having similar cellular localisations (Fig. S2A). To confirm these results we repeated the TRAIL reporter assays in MRC-5 fibroblast cells and also detected reduced transcriptional activity of the phospho-mutants (Fig. 3F). We next generated FLAG-IRF1 phospho-mimetic mutants T181D and S185E and tested their activity on the TRAIL reporter in Cos7 and MRC5 cells. Somewhat surprisingly, these mutants also exhibited reduced activity (Fig. 3F). It should be noted that substitution of aspartate or glutamate can chemically resemble phosphoserine and phospho-threonine residues and thus mimic some functions, the geometry surrounding phosphate group required for proper recognition by binding proteins may not be fully satisfied. However, taken together, we can conclude from our data T181 and S185 residues are required for IRF1 function in cells and that their substitution with alanine or acidic residues is not compatible with full IRF1 transactivation ability.

**Figure 3.**
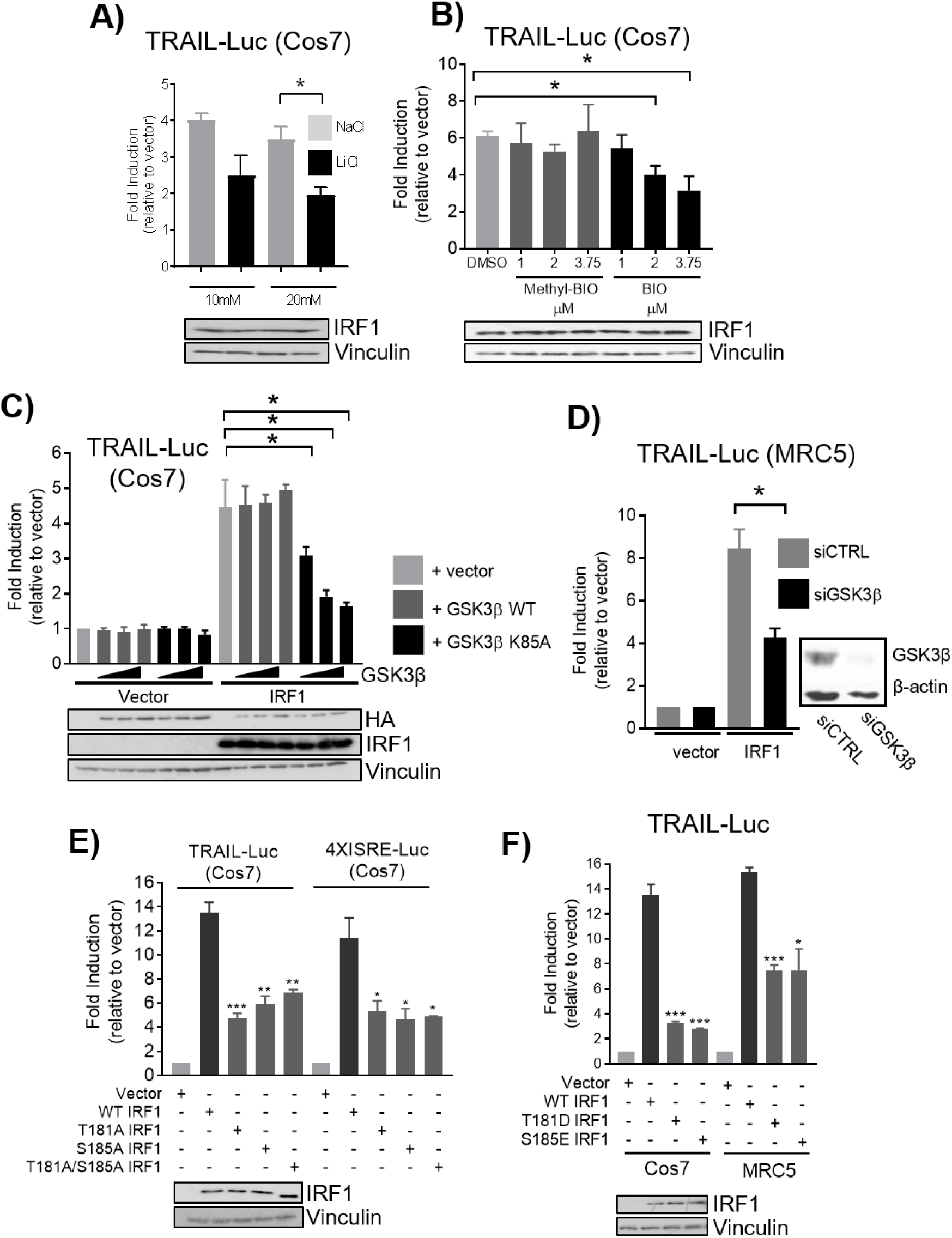
GSK3β is required for IRF1 transcriptional activity. **A**) Reporter assays in Cos7 cells transfected with FLAG-IRF1 and TRAIL promoter reporter for 48 hours. Cells were treated with NaCl (to control for osmolality) or LiCl for 24 hours prior to lysis. Data is expressed as fold induction by IRF1 over vector. All reporter assay data is from three independent experiments assayed in triplicate. Error bars denote SEM and * denotes statistical significance (*p*=<0.05) as determined by Students t-test between NaCl and LiCl treatments. Panel below shows IRF1 expression in lysates. **B**) As for (**A**) but treatment with vehicle (DMSO), GSK3 Inhibitor BIO or the inactive analog Methyl-BIO (1, 2.5 and 3.75µM / 1hr). **C**) Reporter assays using Cos7 cells transfected with TRAIL reporter construct, pcDNA3, or FLAG-IRF1 and increasing concentrations of GSK3β-HA WT or GSK3β-HA K85A mutant. **D**) Reporter assays in MRC5 cells transfected with control or GSK3β siRNAs overnight (10nM) followed by transfection with TRAIL promoter reporter and IRF1 for 24 hours. **E**) Reporter assay in Cos7 and cells transfected with the TRAIL or 4XISRE - Luc reporters in conjunction with FLAG-IRF1 wild type (WT), T181A, S185A, T181A/S185A constructs. Statistical difference is between WT and mutant IRF1. **F**) As for (**E**) but with T181D and S185E mutants in Cos7 and MRC5 cells.

### Expression of IRF1 target genes is dependent on T181 integrity

Having established that GSK3β and the T181 residues are required for IRF1 activity in reporter assays, we next assessed the effects of IRF1 substitution mutations on activation of endogenous gene targets. As the T181A, S185A and T181A/S185A mutants showed indistinguishable activities, we focused on the T181A mutant. Therefore, we generated tetracycline-inducible H3396 cell lines to conditionally express wild-type IRF1 or T181A mutant with an empty vector control. As shown in Fig. S2C, strong induction of IRF1 proteins was observed after 24hrs Dox treatment. We used these cell lines to assess the effect of conditional expression IRF1 WT and T181A proteins on endogenous IRF1 target genes using RT-qPCR. In contrast to the negative control, conditional expression of IRF1 WT protein in H3396 cells resulted in increased expression of TRAIL, OAS3, PSMA6 and TBC1D32 (TBC1 domain 32, previously C6orf170) transcripts. However, induction of these genes by IRF1 T181A was significantly lower (Fig. 4A), consistent with the results from the reporter assays. We next confirmed that the Tet-inducible IRF1 proteins were being recruited to promoters by chromatin immunoprecipitation (Fig. 4B), and found IRF1 T181A was detected at similar or higher levels as IRF1 WT. Thus despite its efficient expression and robust recruitment to IRF1 target gene promoters, these results indicate that T181 integrity is essential for full IRF1 transcriptional activity.

**Figure 4.**
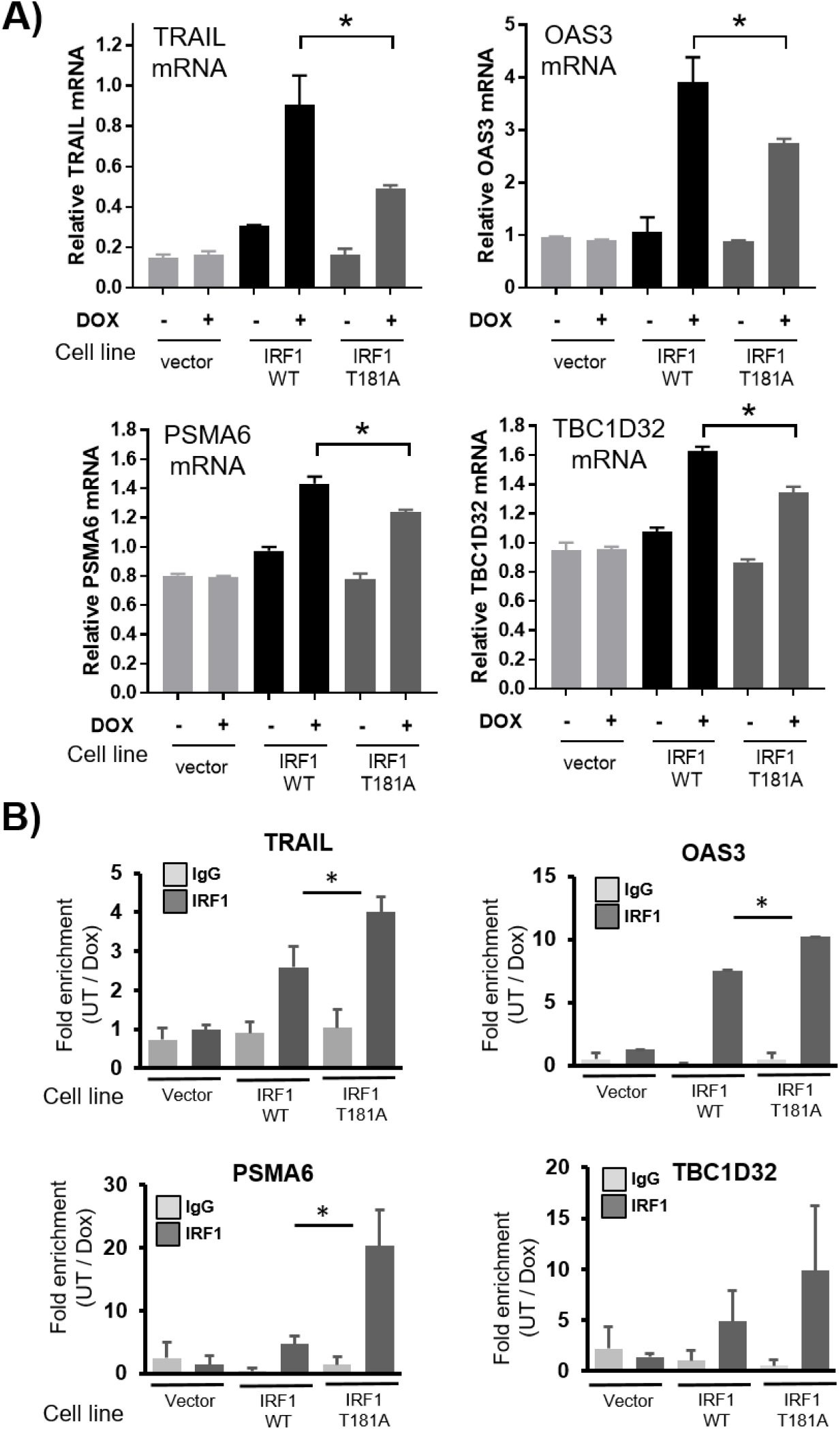
T181 is required for full IRF1 transactivation of target genes. **A**) IRF1 target gene mRNA expression determined by qRT-PCR. H3396-Tet-Off cells expressing empty vector (pCDNA4-TO), WT or T181A IRF1, were induced with 2µg/mL doxycycline for 36hrs. TRAIL, OAS3, PSMA6 and C6orf170 mRNA is expressed relative to β-actin. Statistical significance was determined between Dox-induced WT and T181A. **B**) ChIP analysis performed on the cell lines from (A) using either control rabbit IgG antibody or IRF1 M20 (mouse-specific) antibody to prevent any endogenous IRF1 immunoprecipitation. Data is shown as fold enrichment between cells treated with doxycycline or vehicle.

### IRF1 T181A hampers RNA Pol-II elongation on the C6orf170 gene

Our data indicated that the T181A mutation reduces IRF1 transcriptional activity without compromising its recruitment to target promoters (Fig. 4B). Therefore to further investigate the mechanism underlying the unproductive IRF1 activation due to T181A mutation, we investigated RNA Pol-II phosphorylation status at an IRF1 target gene. The transition of RNA polymerase to an effective transcriptional elongating form is well established (33) and involves phosphorylation events, within the CTD (C-terminal domain) repeat region of Pol-II. We performed ChIP assays on chromatin isolated from H3396 cell lines conditionally expressing WT or T181A IRF1, using antibodies to immunoprecipitate total RNA Pol-II, or elongating Pol-II (phospho-S2) (Fig. 5A/B). We selected the *TBC1D32* gene as its promoter has robust resident Pol-II content in unstimulated cells, thus we reasoned this might facilitate observing the dynamics of Pol-II transition to a transcriptional elongating form (p-Ser^2^), as promoters with a large amount of existing Pol-II are usually enriched in the initiating p-Ser^5^ form of Pol-II CTD [31, 32]. Indeed, a significant proportion of IRF1 targets that we previously identified by ChIP-chip [33] fall into this category. Therefore, we considered the *TBC1D32* gene to be a good candidate to investigate how perturbation of IRF1 phosphorylation might affect the Pol-II transition on IRF1 target genes. As shown in Fig. 5A we observed a reduction in total Pol-II content at the TBC1D32 promoter following Dox stimulation of IRF1 WT expression, but not in IRF1 T181A expressing cells. In the case of IRF1 WT, but not T181A, this was accompanied by an increase in the amount of pSer^2^ Pol II detected within the TBC1D32 gene body, suggesting transition to elongating Pol II. Thus, these data suggest that IRF1 T181 phosphorylation is required for the transition of promoter-bound RNA polymerase II to a transcriptional elongating form for effective expression of an IRF1 target gene.

**Figure 5.**
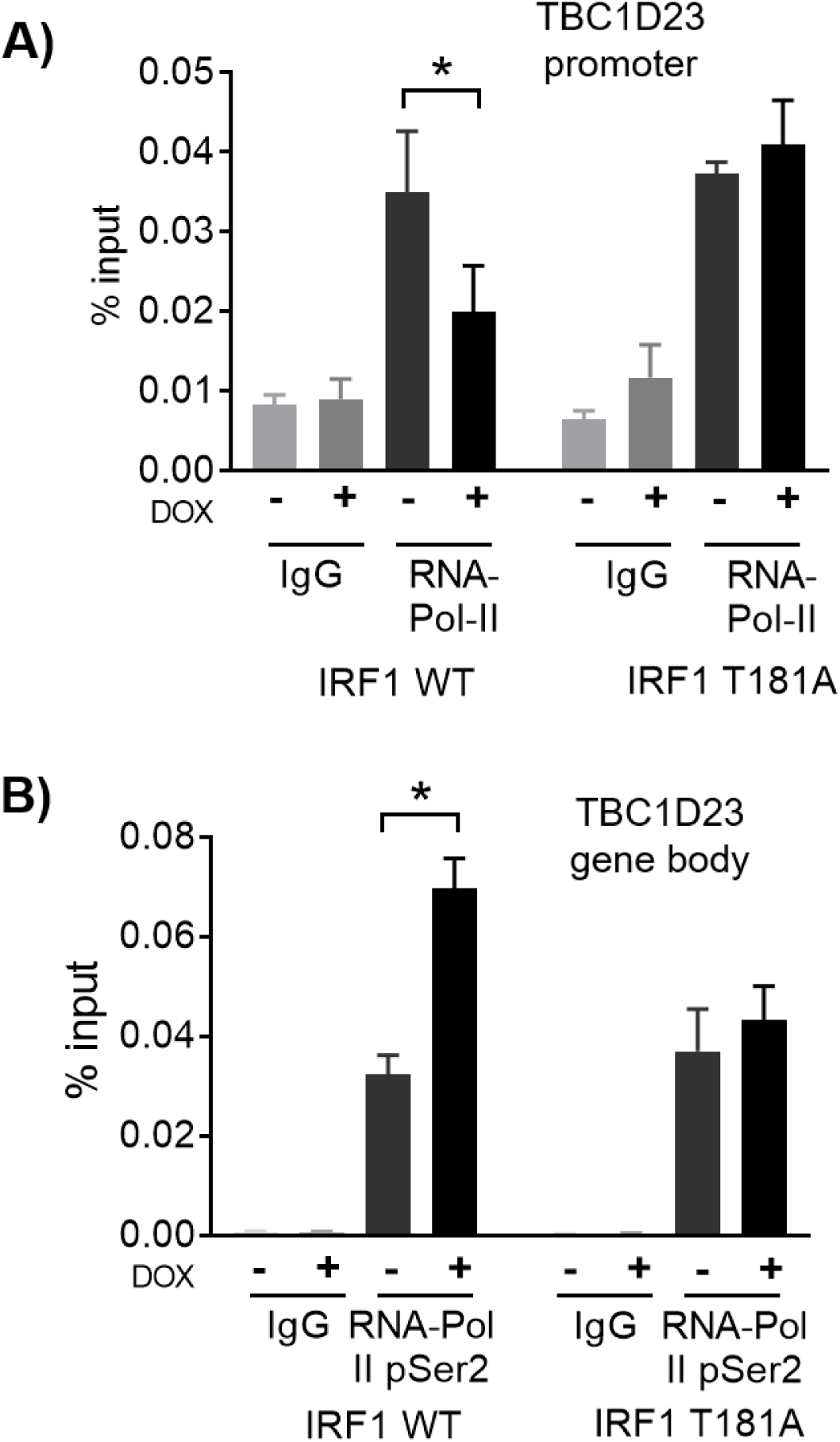
Thr^181^ is required for IRF1 to promote RNA Pol-II elongation on target promoters. **A**) H3396 cells expressing Tet inducible IRF1 WT or T181A proteins were treated with vehicle or induced with Doxycycline for 36 hours. ChIP was performed using anti-total RNA-Pol-II, or IgG antibodies (as control). QPCR was performed to detect enrichment at the C6orf170 promoter region comprising the IRF1 binding site. **B**) ChIP performed as in A, but using anti-phospho-Ser^2^ RNA Pol-II antibody and QPCR performed using primers from within the C6orf160 gene body to detect the elongating form of RNA-Pol-II.

### Phosphorylation of T181/S185 by GSK3β promotes IRF1 degradation

Phosphorylation by GSK3β is known to impact on the ubiquitination and turnover of some of its substrates, e.g. c-Myc and c-Jun, which contain similar sequences to the TPALSP motif in IRF1 (Fig. S1A) [24]. Further we had noted that over-expression of GSK3β resulted in reduced IRF1 (WT) levels (Fig 1E lane 6). Therefore, we tested whether phosphorylation site mutation would alter half-life of IRF1 proteins in MRC5 or HEK293 cells by CHX chase assays. As shown in Fig. 6A&B, T181A, S185A and T181A/S185A mutants displayed decreased turnover, suggesting stabilisation of these proteins compared to WT. In contrast, phospho-mimetic mutations T181D and S185E revealed faster turnover than wild type IRF1. (Fig. 6A & B). Similar results were observed in both MRC5 and HEK293 cells (Fig. S3A). We next assessed the effect of GSK3β overexpression on the half-life of IRF1 WT protein (Fig. 6D-F). Overexpression of WT GSK3β in MRC5 cells strongly reduced IRF1 stability in comparison to the vector only control (Fig. 6C & D). In contrast, overexpression of the kinase inactive GSK3β K85A mutant appeared to increase the stability of IRF1. Similar effects on the half-lives of exogenous IRF1 proteins were observed in HEK293 cells (Fig. S3B). Overexpression of WT GSK3β did not alter the half-life of T181A IRF1 suggesting the destabilisation of IRF1 by GSK3β requires this residue (Fig. S3C). Consistent with these data, depletion of endogenous GSK3β resulted in increased stability of both endogenous (human) and FLAG-tagged (mouse) IRF1 proteins (Fig. 6E & F). Pre-treating cells with GSK3 inhibitor BIO prior to CHX chase also resulted in a stabilisation of IRF1 protein (Fig. S3D). Taken together, these data demonstrate convincingly that phosphorylation of IRF1 at Thr^181^/Ser^185^ regulates its stability.

**Figure 6.**
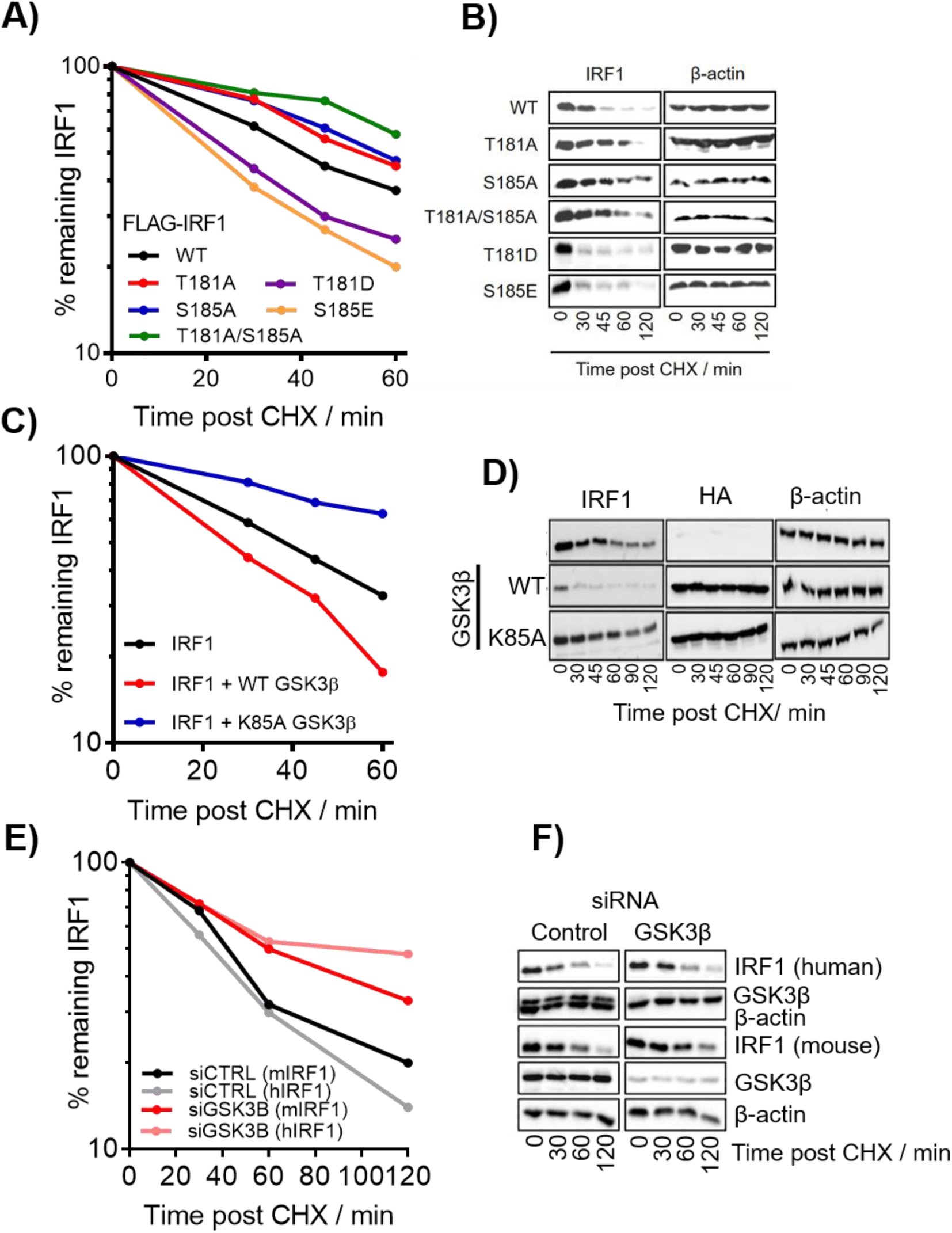
Phosphorylation of Thr^181^/Ser^185^ by GSK3β promotes IRF1 degradation. **A**) Cycloheximide (CHX) chase to detect turnover of IRF1 proteins. MRC5 cells expressing IRF1 WT or indicated mutants were treated with CHX to prevent further protein synthesis. Whole cell extracts were prepared at the times indicated post CHX treatment and subjected to western blotting, using anti-IRF1 antibody. IRF1 expression was quantified using densitometry (ImageJ) and expressed relative to β-actin levels; untreated was set at 100%. Data is from three independent experiments performed in duplicate. Error bars indicate SEM **B**) Western blot of IRF1 CHX chases related to (A). **C**) CHX chase in MRC5 cells expressing IRF1 and GSK3β, calculated as for (A). **D**) Western blot of IRF1 and GSK3β related to (D) **E**) MRC5 cells transfected control or GSK3β siRNAs (10nM) and FLAG-IRF1 for 48hrs and subjected to CHX chase for indicated times. The same lyates were probed with FLAG to detect exogenous IRF1 and IRF1 C20 antibody to detect the endogenous IRF1. Endogenous IRF1 expression was induced by 3hr treatment with 1000U/ml IFNγ prior to CHX chase. **F**) Western blots from E) probed for FLAG (exogenous mouse IRF1) and human IRF1 (C20 antibody is non cross reactive with murine IRF1), GSK3β (knockdown efficiency) and actin loading control.

### GSK3β promotes IRF1 ubiquitination

IRF1 has previously been shown to be poly-ubiquitinated in cells (11) which promotes its degradation and short half-life. We determined the level of ubiquitination on the mutated forms of IRF1 by western analysis of denaturing nickel pulldowns of 6xHis-myc-ubiquitin modified proteins in the presence of MG132. This showed that wild-type IRF1 was robustly ubiquitinated while the alanine mutants were less efficiently ubiquitinated in cells. In contrast, the acidic mutants had increased ubiquitination compared to wild-type IRF1 (Fig.7A/B). There was also an increase in IRF1 ubiquitination upon co-transfection with GSK3β WT, but not GSK3β K85A vectors (Fig. 7C). The requirement of GSK3β for endogenous IRF1 ubiquitination in human cells was determined by siRNA depletion or chemical inhibition. In both cases a large reduction in ubiquitinated IRF1 was detected following ubiquitin IP in H3396 cells (Fig. 7D). Collectively these data support the hypothesis that GSK3β-mediated phosphorylation of T181/S185 is required for regulation of IRF1 turnover via ubiquitination.

**Figure 7.**
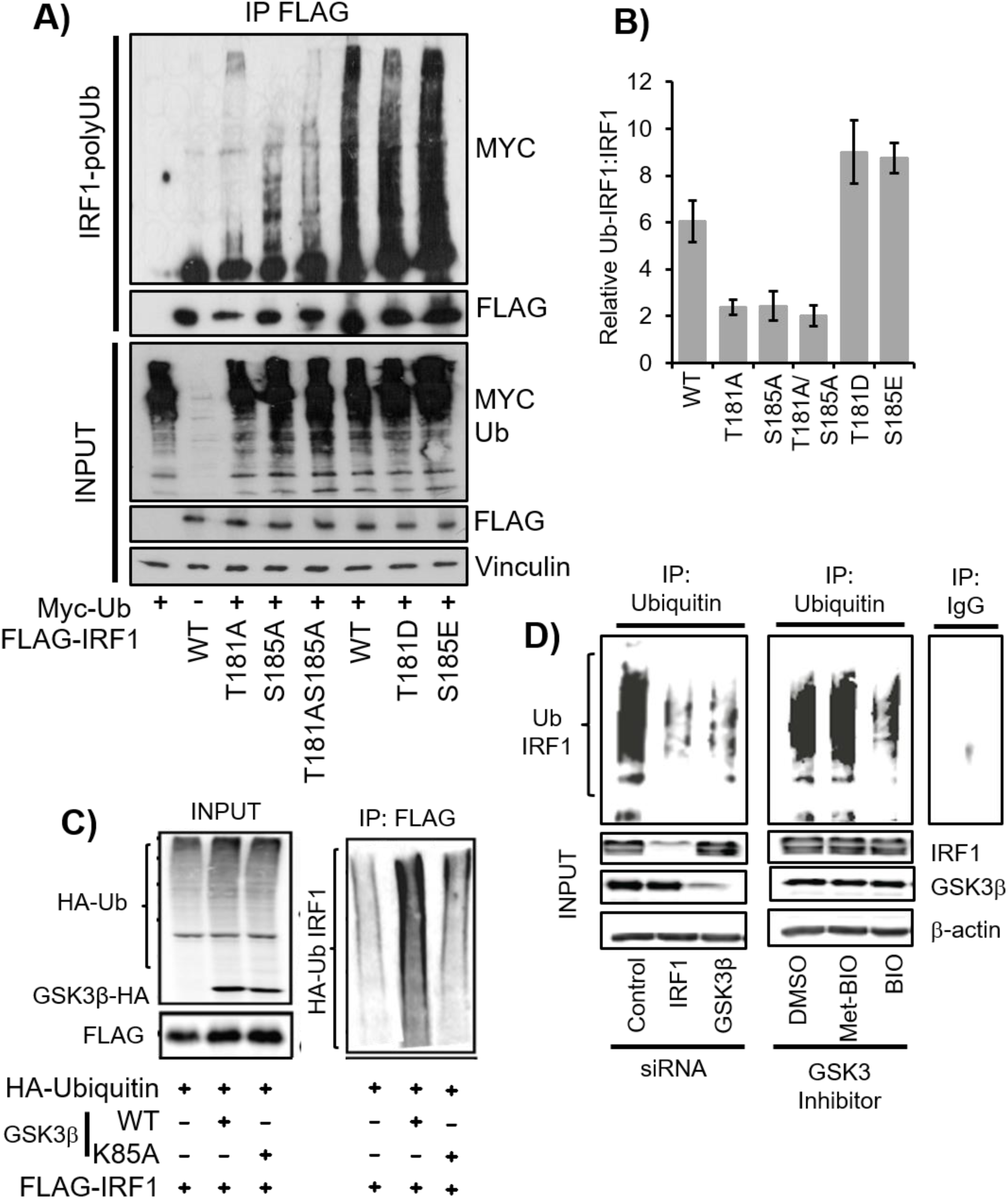
GSK3β and Thr^181^/Ser^185^ promotes IRF1 ubiquitination. **A**) Ubiquitination of IRF1; HEK293 expressing FLAG-IRF1, and 6xHis-myc ubiquitin were treated with MG132 (10uM) for 6 hours prior to lysis and FLAG IP. Lysates were probed with anti-myc and FLAG. **B**) Quantification of relative levels of ubiquitination of IRF1 proteins. Data is expressed as the relative levels of the ubiquitinated species versus the IRF1 from inputs (to account for differences in expression). Data is from 3 experiments. Error bars denote SEM. **C**) HEK293 cells expressing HA-Ubiquitin, FLAG-IRF1 WT, GSK3β-HA WT and GSK3β-HA K85A were lysed 48 hours post transfection in SDS denaturing buffer, boiled and diluted 10x in PBS and immunoprecipitated with FLAG. The resulting high molecular weight ubiquitin modified IRF1 was detected by FLAG western blot. Input panel shows expression of transfected proteins **D**) Ubiquitin immunoprecipitation of endogenous IRF1 in MRC5 lysates from cells siRNA depleted of IRF1 or GSK3β, or pre-treated with GSK3 Inhibitor BIO- or its inactive analog methyl-BIO (10µM for 1 hour). MG132 (10µM for 5 hours) was added prior to lysis to prevent degradation of ubiquitinated IRF1. Ub-IRF1 smears were detected by blot against human IRF1 using the C-20 antibody. Knockdown efficiency of IRF1 is shown in the input panel. Control IgG immunoprecipitation is shown on the adjacent panel.

### IRF1 phosphorylated at Thr^181^/Ser^185^ is linked to transcription and degradation

To determine if the T181/S185-phosphorylated pool of IRF1 is targeted for proteasomal degradation we expressed GSK3β and IRF1 in cells, treated with MG132 and detected phosphorylation as before. We found that MG132 lead to a significant increase in the proportion of IRF1 phosphorylation at Thr^181^/Ser^185^ in both basal and GSK3β overexpression conditions (Fig. 8A). This suggests that this phosphorylated form of IRF1 is targeted for proteasome degradation. We next sought to determine if IRF1 is phosphorylated when bound to DNA by using a DNA binding domain mutant YLP-A (Y109A/L110A/P113A) (34). Lysates of HEK293 cells expressing IRF WT or the IRF1-YLP-A mutant were fractioned into cytoplasmic, soluble nuclear and chromatin pools and probed for IRF1 (Fig. 8B lower panels). Only a small proportion of WT and T181A IRF1 were detected in the cytoplasmic fraction, the DBD mutant YLP-A was more enriched. This was also evident by indirect immunofluorescence (Fig S2A). The chromatin fractions contain roughly equal amounts of WT and T181A, but less IRF1 YLP-A suggesting this fraction contains DNA bound IRF1. After adjusting for IRF1 expression, nuclear and chromatin lysates were immunoprecipitated and probed with pT/S antibody. The WT IRF1 was abundantly phosphorylated in the chromatin-enriched lysates while the YLP-A mutant was poorly phosphorylated (Fig. 8B). This suggests that DNA binding functionality and the association of IRF1 with chromatin facilitate T181/S185 phosphorylation. To determine if T181A stabilises the chromatin-enriched pool of IRF1, we performed CHX chases and separated lysates into soluble and insoluble fractions. While there was no difference in stability between the WT and T181A mutant in the soluble pool, the chromatin pool of T181A was more stable than the WT (Fig. 8C). This confirms that T181 contributes to de-stabilisation of chromatin-associated IRF1. Next we sought to determine if transcriptional elongation regulates IRF1 phosphorylation using the RNA-Pol-II elongation inhibitor DRB (5,6-Dichloro-1-β-D-ribofuranosylbenzimidazole). Indeed, co-immunoprecipitations between IRF1 and GSK3β (Fig 8D) and detection of pT181/S185 (Fig 8E) were both reduced by DRB. Therefore phosphorylation at T181/S185 occurs on DNA bound IRF1, requires RNA-Pol II elongation and promotes proteasomal degradation.

**Figure 8.**
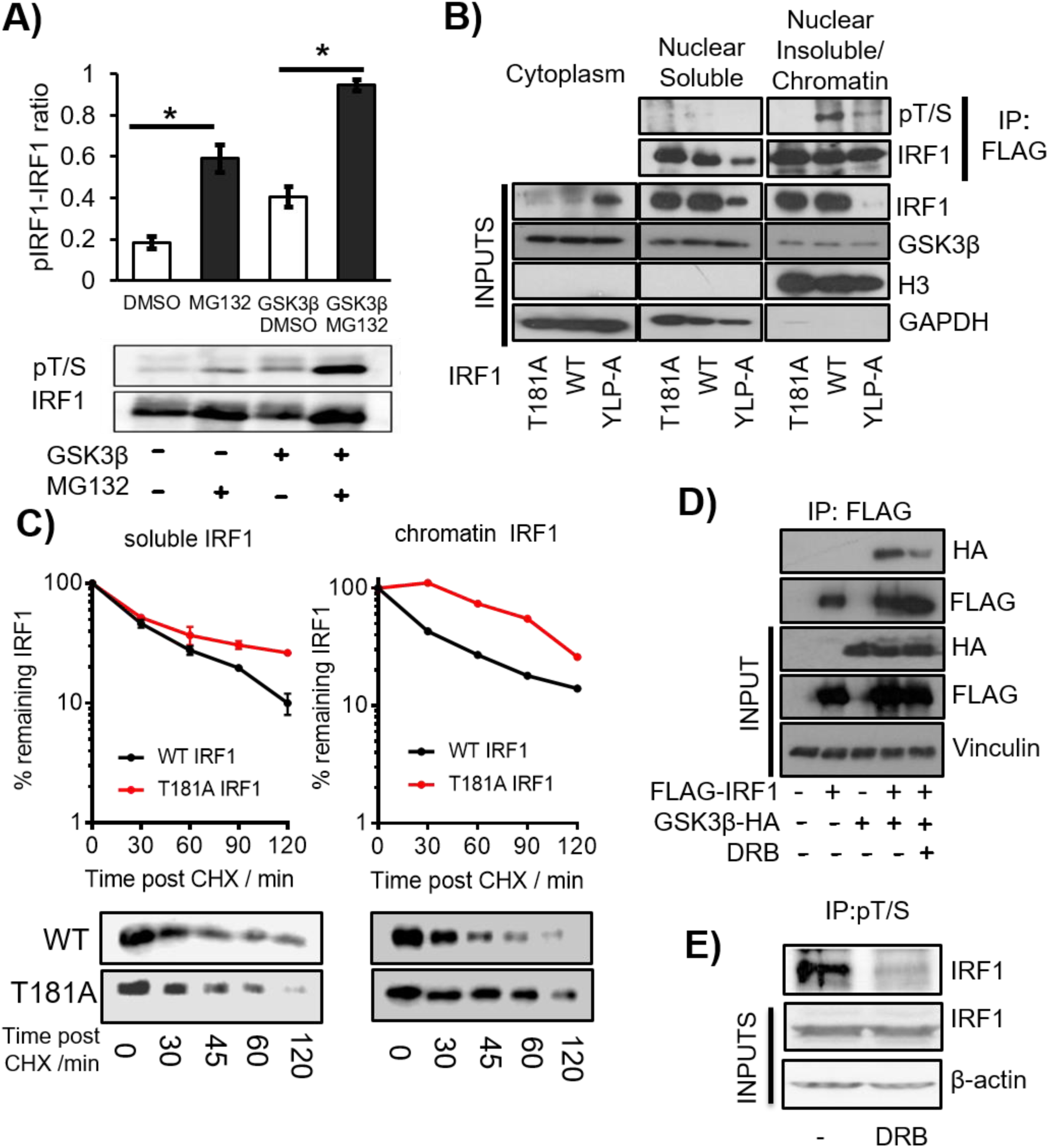
IRF1 phosphorylated at Thr^181^/Ser^185^ is linked to transcription and degradation. **A**) HEK293 cells expressing FLAG-IRF1 with GSK3β-HA or empty vector were treated with 10µM MG132 or DMSO for 6 hours prior to lysis. Following immunoprecipitation with anti-FLAG beads, western blots were performed using the anti-pT/S antibody and re-probed with anti-FLAG antibody to determine total IRF1 protein, and a representative blot is shown. The ratio phospho-IRF1 to total IRF1 was quantified by densitometry, and data from three independent experiments are shown. Error bars denote SEM and * indicates p>0.05 by Students t-test. **B**) Extracts from HEK293 cells expressing WT, T181A or YLP-A IRF1 proteins were separated into cytoplasmic, nuclear and chromatin fractions. Nuclear and chromatin lysates were immunoprecipitated with anti-FLAG after adjustment for IRF1 expression levels and blotted with the anti-pT/S antibody. Lower panels show expression of IRF1 mutants in fractions and GAPDH as a cytoplasmic marker and Histone H3 as a chromatin marker. **C**) HEK293 cells expressing WT or T181A IRF1 were CHX chased for indicated times, lysates were prepared in 200mM NaCl buffer (nuclear soluble) and insoluble pellets (chromatin) were further digested by incubation in 500mM NaCl buffer supplemented with DNase I. **D**) HEK293 cells expressing FLAG-IRF1 and GSK3β-HA were treated with DRB (1μM/ 1hr) prior to lysis and immunoprecipitation with FLAG. **E**) H3396 cells were pre-treated with DRB (1μM/ 1hr) to inhibit transcription prior to immunoprecipitation with IRF1 and blot with pT/S antibody.

### Fbxw7α interacts with IRF1 via Thr^181^

Many GSK3β substrates are targeted for ubiquitination by the SCF E3 ubiquitin ligase receptor Fbxw7, and we noted that phosphorylation of T181/S185 would generate an Fbxw7 phosphodegron. We tested IRF1 interaction with various isoforms of Fbxw7 (α/β/γ). HEK293 lysates expressing GST-Fbxw7 isoforms were subjected to GST pulldowns and demonstrated that IRF1 preferentially interacts with the nuclear form; Fbxw7α (Fig. S4A/B). The interaction was confirmed by reciprocal co-IP in HEK293 cells (Fig. 9A/B). IRF1 did not interact with a WD40 domain deletion of Fbxw7α (Fig. 9B), suggesting the interaction with IRF1 is phosphorylation dependent as observed for other substrates. The interaction was substantially increased when IRF1 phosphorylation was enhanced by GSK3β over-expression (Fig. 9C). Further, IRF1 T181D increased, while T181A decreased interaction with Fbxw7α (Fig. 9D). These data demonstrate a phosphorylation dependent interaction between IRF1 and Fbxw7α.

**Figure 9.**
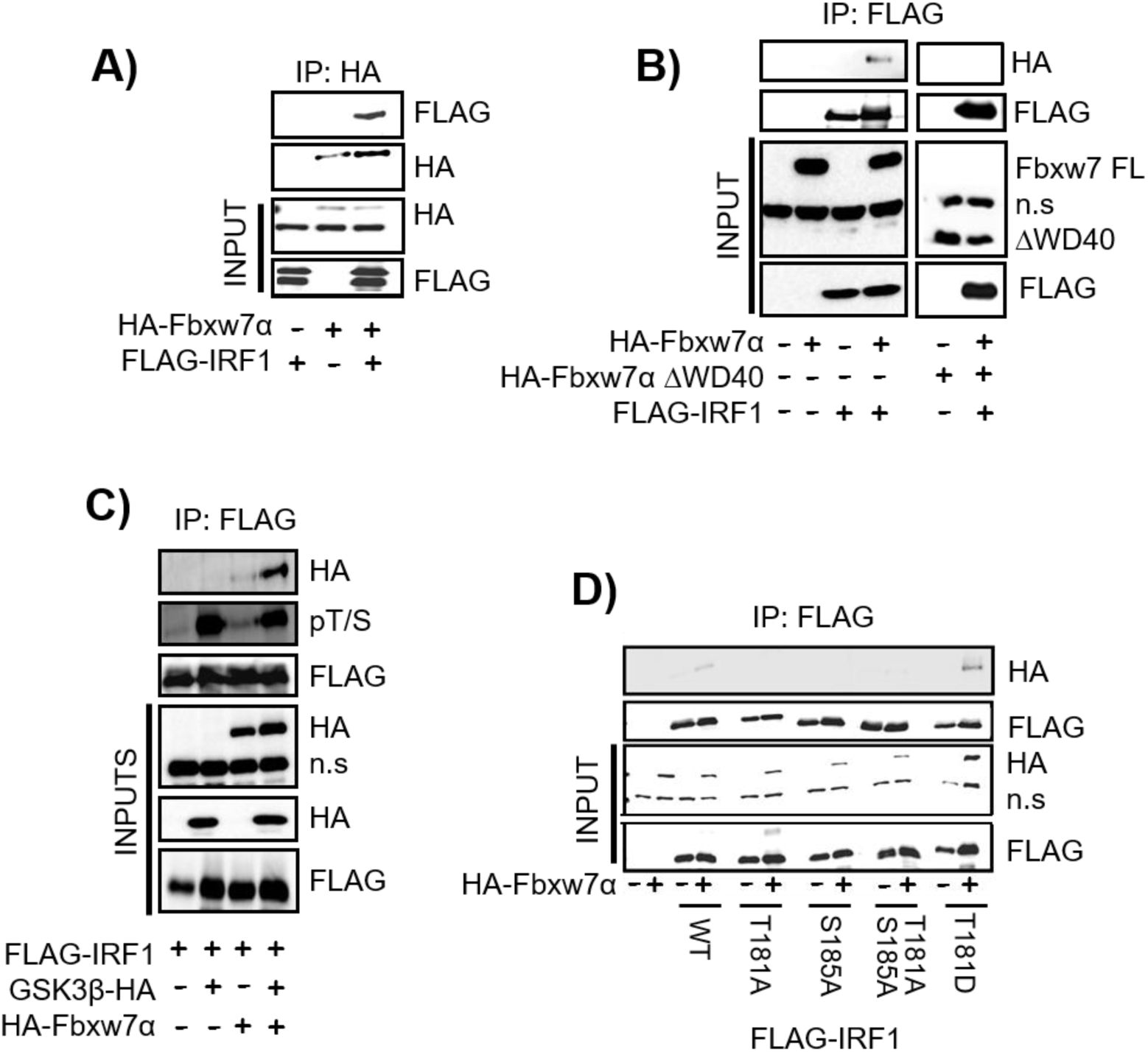
Fbxw7α association with IRF1 requires T181 integrity. **A**) Co-immunoprecipitation of HA-Fbxw7α in extracts of HEK293 cells, and western blots using anti-FLAG antibody to detect associated FLAG-IRF1. Cells were treated with MG132 for 6 hours prior to lysis. Blots were re-probed with anti-HA to determine IP efficiency. N.s. = non-specific band is indicated. **B**) Co-IP experiment as in (**A**) but using with anti-FLAG antibody to IP and western blot with anti-HA to detect complex formation with HA-Fbxw7α or a mutant lacking the WD40 domain (HA- Fbxw7α ΔWD40). **C**) HA-Fbxw7α, GSK3β-HA and FLAG-IRF1 were co-expressed in HEK293 for 48 hours, 6 hours prior to lysis cells were treated with 10µM MG132. Lysates were immunoprecipitated with anti-FLAG and probed with anti-HA to detect Fbxw7α interaction. The blots were also probed with the anti-pT/S antibody and FLAG to determine relative levels of IRF1 phosphorylation. **D**) Immunoprecipitations as in (**C**) but with the inclusion of IRF1 T181A, S185A, T181A/S185A and T181D mutants.

### Fbxw7α regulates IRF1 ubiquitination, half-life and transcriptional activity

We next determined what effect Fbxw7α has on the stability of IRF1. Co-transfection of Fbxw7α with IRF1 resulted in decreased stability of IRF1 in MRC5 CHX chase experiments. No effect on IRF1 half-life was detected upon co-transfection with ΔWD40 Fbxw7α (which does not interact with IRF1) (Fig. S5A / B). Conversely, Fbxw7 siRNA increased the half-life of IRF1 (for both exogenous FLAG IRF1 and endogenous human IRF1) in MRC5 (Fig. 10A & S5C). We also performed His-Ub^WT^ and Ub^K48O^ (K48 only) pulldowns of IRF1 with GSK3β or Fbxw7α overexpression and detected an increase in IRF1 ubiquitination when either of these proteins were overexpressed (Fig. 10B). This suggests GSK3/Fbxw7 promotes K48-linked ubiquitination of IRF1. To determine if this ubiquitination was important for IRF1 function we measured IRF1 activity in Fbxw7 depleted cells and noted a significant reduction in TRAIL reporter activity (Fig. 10C). Next we wanted to further confirm the importance of Fbxw7 in IRF1 function so sought to map the lysine residues targeted by the SCF^Fbxw7^ ligase. As GSK3β-Fbxw7α targets DNA bound IRF1 we reasoned that lysines within the DBD might be masked, and thus focussed on the remaining five C terminal lysine residues. Each of the K→R mutants were expressed with and without Fbxw7 and subjected to His-Ub pulldowns as before. Increased ubiquitination following Fbxw7 overexpression was observed with the WT, K276R and K300R variants, while K233R, K240R and K255R were less efficiently hyper ubiquitinated, suggesting these residues, and in particular K240 may serve as targets of SCF^Fbxw7α^ (Fig. 10D & S5D). These mutants also show reduced overall ubiquitination with the Ub^WT^ and Ub^K48O^ mutant but not the Ub^K63O^ or Ub^K6O^ variants suggesting they are acceptors of K48-Ub linkages, while K6 and K63-Ub chains are formed on other lysine residues (Fig. S5E). The Fbxw7α insensitive mutants are also more stable (Fig. 10E-F) and less transcriptionally active on both the TRAIL and 4XISRE reporters (Fig. 10G & S5F). Collectively, these data demonstrate that Fbxw7α ubiquitinates DNA bound IRF1 at several residues with the TAD region which regulates its stability and transcriptional activity.

**Figure 10.**
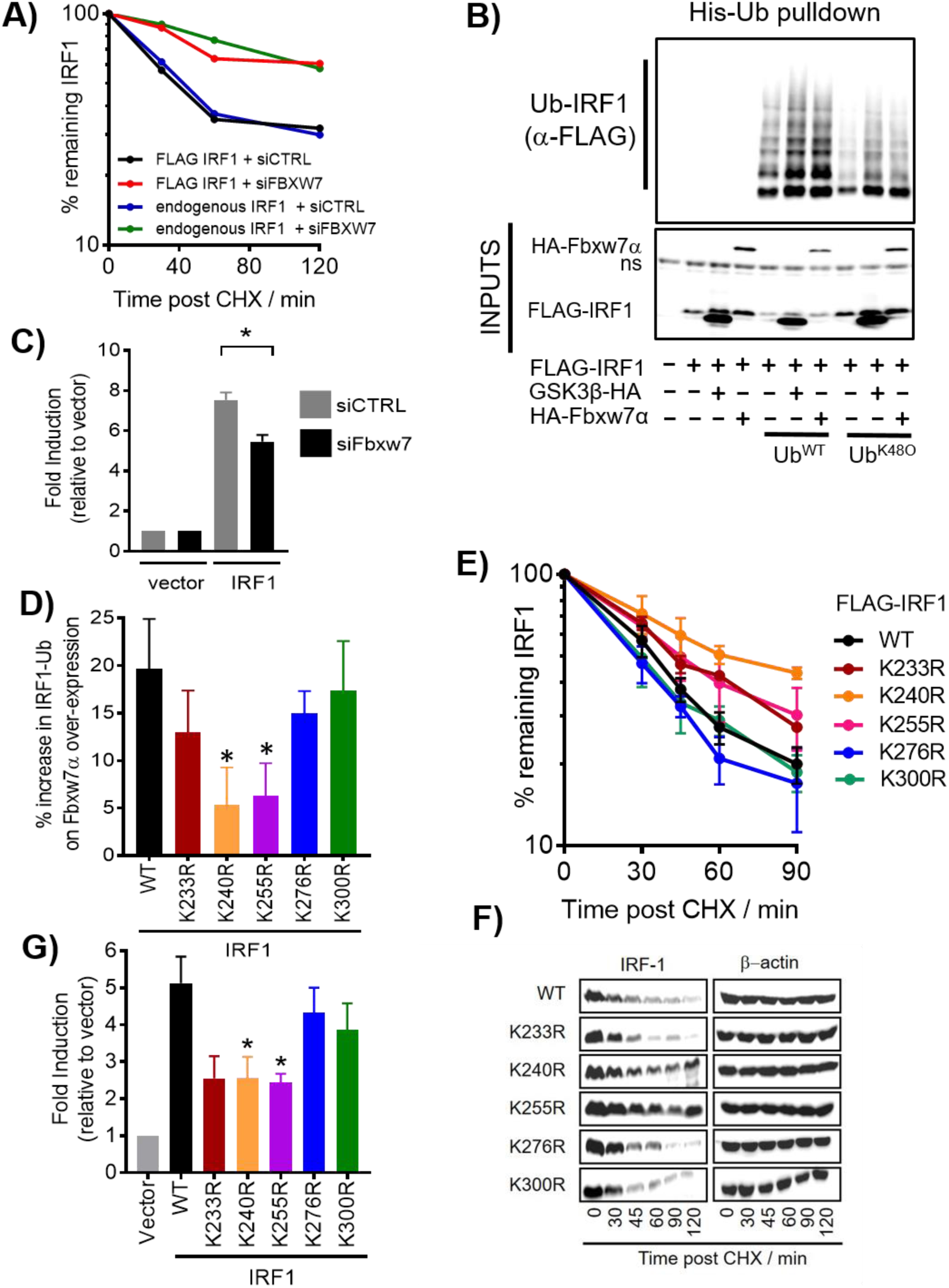
Fbxw7α regulates IRF1 ubiquitination, half-life and transcriptional activity. **A**) CHX chase of MRC5 cells expressing IRF1, HA-Fbxw7α, HA-Fbxw7α ΔWD40 mutant. Percentage remaining IRF1 was calculated relative to untreated control, and normalised to the loading control β-actin. Both exogenous (FLAG) and endogenous IRF1 were assayed from the same lysates. **B**) HEK293 cells expressing FLAG-IRF1, GSK3β-HA and 6xHis-Ub (WT or K48 only) for 48 hours prior to a 6 hour 10µM MG132 treatment. Lysates were prepared in 8M urea buffer and incubated with nickel agarose to enrich His-Ub modified proteins. Pulldowns were probed with anti-FLAG to detect Ub-IRF1. 10% inputs show expression of transfected proteins. **C**) Reporter assay in MRC-5 cells transfected with the TRAIL promoter-Luciferase reporter. Cells were transfected with the indicated siRNAs overnight prior to transfection with reporter and IRF1 expression vector for 24 hours prior to lysis. Error bar denotes SEM and * statistical significance *p*(>0.05) as determined by Students t-test between control and Fbxw7 siRNA treated cells. **D**) Quantification of relative ubiquitination of indicated IRF1 K→R mutants. HEK293 cells expressing FLAG-IRF1, HA-Fbxw7α and 6xHis-Ub WT) for 48 hours prior to a 6 hour 10µM MG132 treatment. Lysates were prepared in 8M urea buffer and incubated with nickel agarose to enrich His-Ub modified proteins. Pulldowns were probed with anti-FLAG to detect Ub-IRF1. Data is shown as % increase in ubiquitination between empty vector and HA-Fbxw7 expressing pulldowns. **E**) CHX chase of MRC5 transfected with indicated IRF1 mutants. **F**) Western blot related to (E)**. G**) Reporter assay in MRC-5 cells transfected with the TRAIL promoter-Luciferase reporter. Cells were transfected with the indicated IRF1 expression vector for 48 hours prior to lysis. Error bar denotes SEM and * statistical significance *p*(>0.05) as determined by Students t-test.

### IRF1 T181 is required for anti-proliferative activity in cancer cells

The anti-proliferative action of IRF1 is well studied and contributes to tumour suppressor activity (3). To investigate the contribution of T181 phosphorylation to IRF1 activity we measured proliferation of H3396 cells using the KRAB-Tet system. As even low IRF1 expression affects cancer cell growth we used this system as it promotes a much more robust silencing of IRF1 expression prior to the addition of doxycycline. We measured proliferation rates in vector, WT and T181A expressing H3396 stable cell lines. Induction of WT IRF1 resulted in a marked decrease in growth of H3396. In contrast, the T181A mutant showed no inhibition of growth similar to empty vector control (Fig. 11A). We next assayed clonal growth in a number of puromycin-selected cell lines constitutively expressing vector, WT or T181A IRF1. Compared to empty vector, WT IRF1 expression reduced clonal growth ability in the majority of lines, while the T181A mutant had less impact. (Fig. 11B). We expanded the analysis with more cell lines and assayed short-term growth effects in puromycin selected cells. As with clonal growth, the number of viable cells was significantly reduced in WT versus T181A expressing lines. (Fig. 11C). A number of cell lines have been identified to harbour deleterious mutations in Fbxw7, these mutations cluster in hotspots within the WD40 repeats and result in a loss of phospho-specific binding [28], additionally some cell lines lack Fbxw7 expression due to homozygous deletions. We tested 7 such lines for their ability to respond to IRF1 overexpression (Fig. 11C & S6A). Remarkably these lines were resistant to the effects of IRF1 expression, suggesting loss of Fbxw7 function may render cancers resistant to some of the tumour suppressor activities of IRF1. We noted that enforced expression of IRF1 did not increase the proportion of non-viable/dead cells suggesting the effect on cell numbers is caused by reduced proliferation rather than increased death. We measured proliferation in vector, WT and T181A cells by scoring the proliferative marker Ki67 and noted a reduction in proliferation in WT IRF1 expressing cells but less so in T181A expressing cells, as before the proliferation of Fbxw7 defective lines were only marginally affected by IRF1 expression (Fig. 11D). Collectively this suggests the T181 residue is essential for the anti-proliferative phenotype of IRF1 expression and that Fbxw7 status is important for this activity.

**Figure 11.**
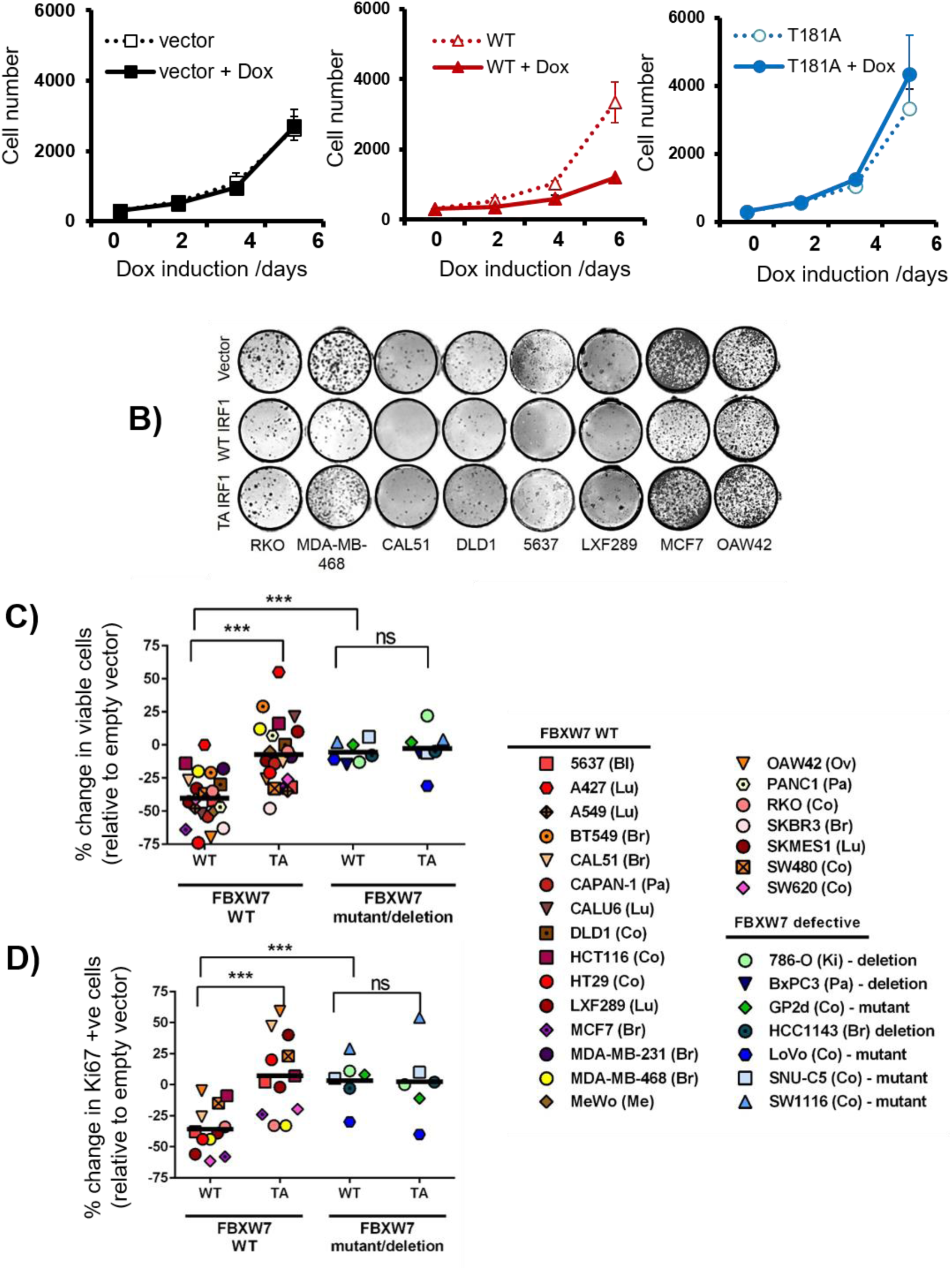
T181 is required for IRF1 anti-proliferative activity in cancer cells. **A**) H3396 stable inducible cell lines expressing IRF1, T181A or vector only were plated at equal concentrations and treated with Dox or vehicle. Proliferation was monitored for 6 days using cell counting. Graph shows the average of three independent experiments performed in quadruplicate +/-standard deviation. **B**) Indicated cell lines were transduced with pBABE-puro, IRF1 WT or IRF1 T181A, selected with puromycin for 48 hours to remove uninfected cells and plated at 500 cells/well on 48-well plates in quadruplicate. Clones were left to grow for 10 days before crystal violet staining. Representative wells are shown. **C**) Cell lines transduced with retroviruses as for **B**) and allowed to grow in puromycin supplemented media for 7 days prior to trypan blue counting. Viable cells were counted from triplicate wells and the % change in viable cell number was calculated relative to empty vector. *** indicates a *p* value less than 0.001 between groups. Abbreviations, Bl (Bladder), Lu (Lung), Br (Breast), Pa (Pancreas), Co (Colorectal), Me (Melanoma), Ov (Ovarian), Ki (Kidney). **D**) Cells treated as for **C)** but plated on coverslips and assayed by indirect immunofluorescence for Ki67 expression. Cells stained with strong nucleolar Ki67 were counted as proliferative. Proliferation was measured relative to vector only transduced cells. Negative values indicate reduction in Ki67/proliferative cells.

## DISCUSSION

Here we report a novel mechanism for IRF1 transcriptional control centring on phosphorylation-dependent degradation. This tightly controlled clearance is important for IRF1 dependent RNA-Pol-II elongation, mRNA generation and anti-proliferative activity.

Several reports have identified ubiquitin E3 ligases that target IRF1. The first identified IRF1 E3 ligase, CHIP ubiquitinated IRF1 using both K48 and K63 linkages (12). Subsequently, MDM2 was found to mono-ubiquitinate IRF1 but did not regulate its turnover (13). The third ligase cIAP2 specifically modifies IRF1 with K63-linked ubiquitin during IL-1 signalling and also does not regulate turnover (15). Each E3 ligase conjugates ubiquitin to different lysine residues, which mostly reside in the DNA binding domain. Ubiquitination site profiling also identified residues within the DBD as the major modified sites (13,15,35). In the case of CHIP, IRF1 is ubiquitinated in the DBD when in its non-DNA bound form. DNA binding thus shields these residues from CHIP dependent degradation (12).

We propose the following model for how ubiquitination and degradation are required for IRF1 transcriptional activity (Fig 12). *De novo* IRF1 protein is induced following stimulation with agents such as IFNγ. The steady state levels of non-DNA bound IRF1 are maintained by E3 ligases (such as CHIP), which target this pool for degradation. IFNγ signalling induces a widespread recruitment of IRF1 to target genes. DNA-bound IRF1 is then protected from degradation as lysines in the DBD are shielded from Ub E3 ligases. The difference in half-life of nucleoplasmic vs DNA engaged IRF1 most likely reflects the ‘time’ required for other signalling/remodelling/recruiting events to occur prior to successful transcriptional elongation. We propose that following firing of RNA polymerase II into the elongation phase, GSK3β dependent phosphorylation marks IRF1 as “spent”, which then targets SCF^Fbxw7^ to ubiquitinate C terminal lysine residues and promote proteasome dependent degradation. Essentially, this represents a transcriptional time clock (36,37). This coordinated response, allows IRF1 to successfully engage the transcriptional apparatus before it is marked for degradation. Importantly, further rounds of transcription would only occur if there is a continued presence of stimuli (e.g. IFNγ) that would produce additional IRF1 protein. This modulation of IRF1 activity likely acts as a restraint to prevent hyper-activation of target genes which includes regulators of processes that need to be tightly controlled-such as inflammation, apoptosis and cell cycle. Interestingly, this regulation must be balanced, as the transcriptional regulation by IRF1 is dually sensitive to perturbation, hypo-or hyper- stimulation (such as the acidic mimic mutants) leads to disruption of IRF1 activity. This suggests that any de-coupling of IRF1 activity from ‘inducing stimuli’ will lead to a restriction of target gene activation.

**Figure 12.**
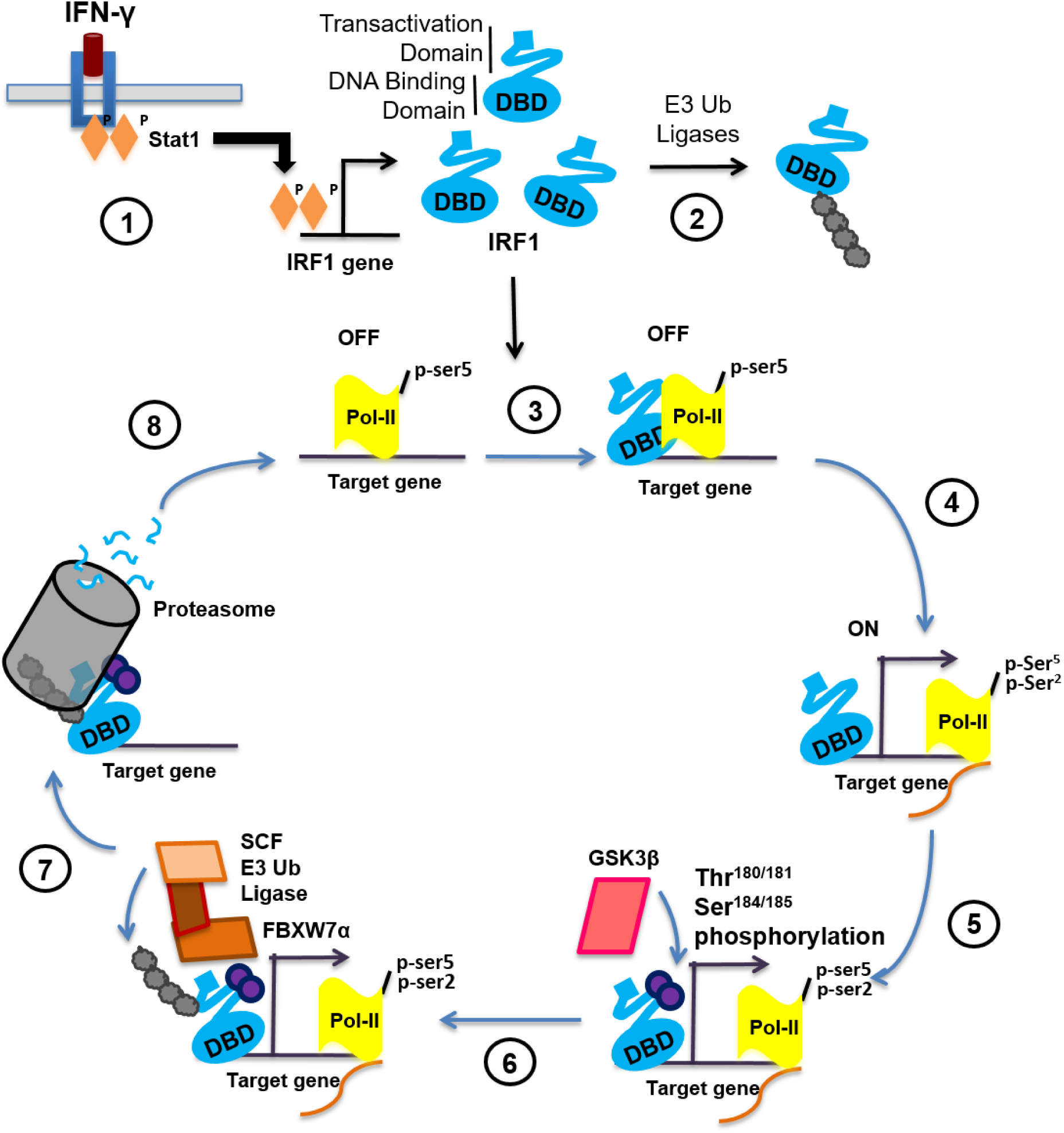
Proposed mechanism of GSK3β-dependent regulation of IRF1 activity. Schematic representation of a proposed model for IRF1 regulation by GSK3 kinases. 1) External stimuli (such as IFNγ signalling activate STAT1 leading to increase expression of cellular IRF1 protein. 2) Steady state levels of IRF1 protein are maintained by the ubiquitin proteasome system. Non- DNA bound IRF1 is ubiquitinated at lysine residues exposed within the DBD. 3) Nuclear IRF1 binds recognition sequences in target promoters. In many cases such as TBC1D32 gene, these IRF1-bound promoters are marked by high levels of pSer5 (initiating) modified RNA-Pol-II and are thus poised for transcription. Engagement with DNA shields the lysines within the DBD from recognition by E3 ligases and subsequent degradation by the proteasome. This allows time for further events that are necessary for transcription to occur. 4) Transcription is initiated, RNA-Pol-II is marked with pSer2 (elongating). 5) Successful initiation triggers phosphorylation of IRF1 at T181/S185 by GSK3β. It is not known how this phosphorylation senses RNA-Poll firing, perhaps a reorganisation of proteins at the promoter un-masks epitopes in IRF1 allowing binding and phosphorylation. 6) Phosphorylation of IRF1 generates a phospho-degron recognised by SCF-Fbxw7, which promotes K48 linked ubiquitination of IRF1. 7) IRF1 is degraded by the proteasome. It is not known if the degradation occurs while IRF1 is bound to DNA, or if a release of IRF1 occurs beforehand. 8) The previously occupied element is now free for additional molecules of IRF1 (or other proteins) to re-bind and begin a new cycle of transcription. Local concentrations of IRF1 protein – determined by the balance between *de novo* generation of IRF1 and degradation will help dictate whether this additional cycle occurs.

We have demonstrated with the T181A mutant that perturbation of this clearance has significant effects on IRF1 function. Failure to efficiently remove IRF1 from promoters appears to prevent further elongation of RNA-Pol II, reduces target gene mRNA transcript abundance and ablates IRF1’s anti-proliferative activity in a number of cell types. The alanine mutants do however retain some transcriptional activity in reporter assays and mRNA induction. While these mutants are more stable than WT IRF1 they are still degraded - perhaps suggesting other pathways can promote IRF1 clearance from promoters.

The dysfunctional nature of T181A may also offer insight into mechanisms that disrupt IRF1 activity. Inability to clear DNA bound IRF1 may also expose it to modification by SUMO (Small Ubiquitin-related Modifier). SUMOylation of C-terminal lysines has been shown to stabilise IRF1 by competing for ubiquitination. Hyper-SUMOylation of IRF1 has been detected in ovarian cancers, and is known to disrupt IRF1 transcriptional activity (38).

Loss of function in the GSK3β-Fbxw7 axis occurs in several cancer types (30). This is recognised as cancer promoting due to the increased abundance of several oncogenic factors such as c-MYC, c-JUN, cyclin E and NOTCH. Indeed the majority of Fbxw7 substrates are oncogenic, which makes IRF1 (a putative tumour suppressor) and unusual substrate for this pathway. Out data suggests that rather than targeting IRF1 to reduce overall abundance – which would impede its tumour suppressive function, GSK3β-Fbxw7α aid in the timely clearance of DNA bound IRF1 to support additional rounds of transcription and thus downstream phenotypes such as reduced proliferation. Indeed cancer cell lines with detective Fbxw7 are largely resistant to the anti-proliferative activities of IRF1 suggesting that Fbxw7 is an important co-activator of IRF1 function. Collectively, our data supports an essential contribution of GSK3β-Fbxw7α to the tightly regulated turnover of IRF1 protein during the transcriptional cycle and, thus its anti-cancer activities (34,39).

## Supporting information

## ACKNOWLEDGEMENT

This work was supported by BBSRC New Investigator grant (BB/F000340/1) and Cancer Research UK grant (C24478/A9004) to NMC. AJG was supported by a PhD stipend from the School of Pharmacy. AJG, LB and JRM were supported by grants from Breast Cancer Campaign (2006NovPHD13), CRUK (C8820/A19062) and the Wellcome Trust (Z06343/Z/17/Z). We are grateful to the School of Pharmacy for equipment and facilities support provided to the Gene Regulation & RNA Biology group. We thank Debbie Evens, Barbara Rampersad and Hilary Collins for technical/ research support. We thank Dr Abdolrahman Shams Nateri for useful comments on the manuscript. We acknowledge the kind gifts of expression vectors used in this study from the following labs; Harper, JW (GST-Fbxw7 plasmids); Prives, C (HA Ubiquitin); Woodgett, J (GSK3β plasmids); Stein, GS (4xISRE-LUC).

## Author Contributions

AJG designed and performed experiments, analysed data, generated stable cell lines and co-wrote the paper. NMC designed and performed experiments, analysed data and edited the manuscript. AR performed ChIP and qPCR experiments; JX performed GST pulldown; LB generated constructs; AHAK performed ubiquitin assays; JRM provided reagents and edited the paper. DMH provided reagents and constructs, performed data analysis and edited the paper. NMC and DMH co-supervised the postgraduate researchers.

