## Supplementary material for "GSK3β-SCF^FBXW7^ mediated phosphorylation and ubiquitination of IRF1 are required for its transcription-dependent turnover"

**Supplemental Figure Legends**

**Supplementary Figure 1. IRF1 is phosphorylated by GSK3β**

**A)** Alignment of the IRF1 GSK3 consensus site with a subset of other known GSK3 substrates, all sequences are human and the location indicated in parenthesis. Grey highlights indicate phosphorylated residues
**B)** Schematic representation of IRF1 functional domains, indicating secondary structure within the DNA binding domain. The TAD (Transcriptional Activation Domain) and Enhancer domains are predicted to be unstructured. The location of the TPALSP motif within the transactivation domain is indicated. The conserved Thr180/181 and Ser184/185 (mouse/ human) GSK3β phosphorylation sites are highlighted in red.
**C)** Immunoprecipitation experiments revealing phosphorylation of IRF1 T181 by GSK3β. HEK293 were transfected with expression vectors for YFP-IRF1 WT, YFP-IRF T181A, YFP-IRF1 S185A or YFP-IRF1 T181A/S185A together with GSK3β-HA vector or empty vector. Whole cell extracts were immunoprecipitated with anti-pT/S antibody and probed with anti-GFP antibody to detect YFP-IRF1 proteins, indicating T181 phosphorylation levels. The middle panel shows detection of YFP-IRF1 proteins and β-actin loading control (at 10% input for IP). The bottom panel shows GSK3β

**Supplementary Figure 2. GSK3β is required for IRF1 transcriptional activity**

**A)** Indirect immunofluorescence of FLAG-IRF1 and mutants in Cos7, scale bar = 10μm
**B)** Reporter assay in MRC5 using the TRAIL-Luc reporter. Statistical differences are determined between WT and mutant IRF1.
**C)** H3393 stable cells lines (vector, WT or T181A IRF1) treated with Dox for either 24 or 36 hours to induce expression of IRF1, β-actin is shown as loading control.

**Supplementary Figure 3. Stability of IRF1 T181A is not affected by GSK3β over-expression and GSK3β inhibition stabilises IRF1**

**A)** Half-lives of IRF1 protein expressed in MRC5 or HEK293 cells determined by CHX assays related to Fig 6A and 6B
**B)** Half-lives in MRC5 and HEK293 related to Fig 6C and 6D
**C)** CHX chase in MRC5 cells related to figure 6D but with the addition of the T181A mutant
**D)** CHX chase in MRC5 cells (expressing WT FLAG-IRF1) pre-treated with the GSK3 inhibitor X (2.5μM) for 1 hour prior to CHX addition and chase as before.

**Supplementary Figure 4. Fbxw7α interacts with IRF1**

**A)** HEK293 transfected with GST or GST-Fbxw7 (α, β, γ) expression plasmids and FLAG-IRF1 for 48 hours. 6 hours prior to lysis cells were treated with MG132 (10μM). Lysates were incubated with Glutathione-Sepharose beads for 3 hours. Captured proteins were revealed by immunoblot with anti-GST and anti-FLAG antibodies. 10% inputs demonstrate expression of transfected proteins.
**B)** Indirect immunofluorescence of GSK3β and FBXW7 in Cos7. Scale bar = 10μm.

**Supplementary Figure 5. Fbxw7α regulates IRF1 ubiquitination, half-life and transcriptional activity**

- **A)** HEK293 CHX chase of cells expressing FLAG-IRF1, HA-Fbxw7α FL or HA-Fbxw7α ΔWD40.
- **B)** Lysates related to S5A, note the ΔWD40 mutant migrates below the nonspecific band detected by the HA antibody.
- **C)** Western blot panel related to figure 10A
- **D)** Ubiquitination of IRF1 K→R mutants in HEK293 co-transfected with Fbxw7. Lysates were enriched for ubiquitinated proteins by Ni2+ pulldown and probed with FLAG antibody. Inputs show expression of transfected proteins.
- **E)** Quantification of ubiquitination status of IRF1 K→R mutants using indicated ubiquitin variants. MRC5 transfected with indicated IRF1 and Ub mutants were assayed as before. Data is for three independent repeats. .
- **F)** Luciferase reporter assays in MRC5 cells expressing indicated IRF1 construct and the 4XISRE-Luc reporter construct.

**Supplementary Figure 6. Thr$^{181}$ and Fbxw7 are required for IRF1 anti-proliferative activity in cancer cells.**

- **A)** Bar graph of data presented in 11C. Data is shown as % change in cell number where empty vector is expressed as 100%. Error bars = SEM.

**Supplemental Figures**

### Figure S1. IRF1 is phosphorylated by GSK3β

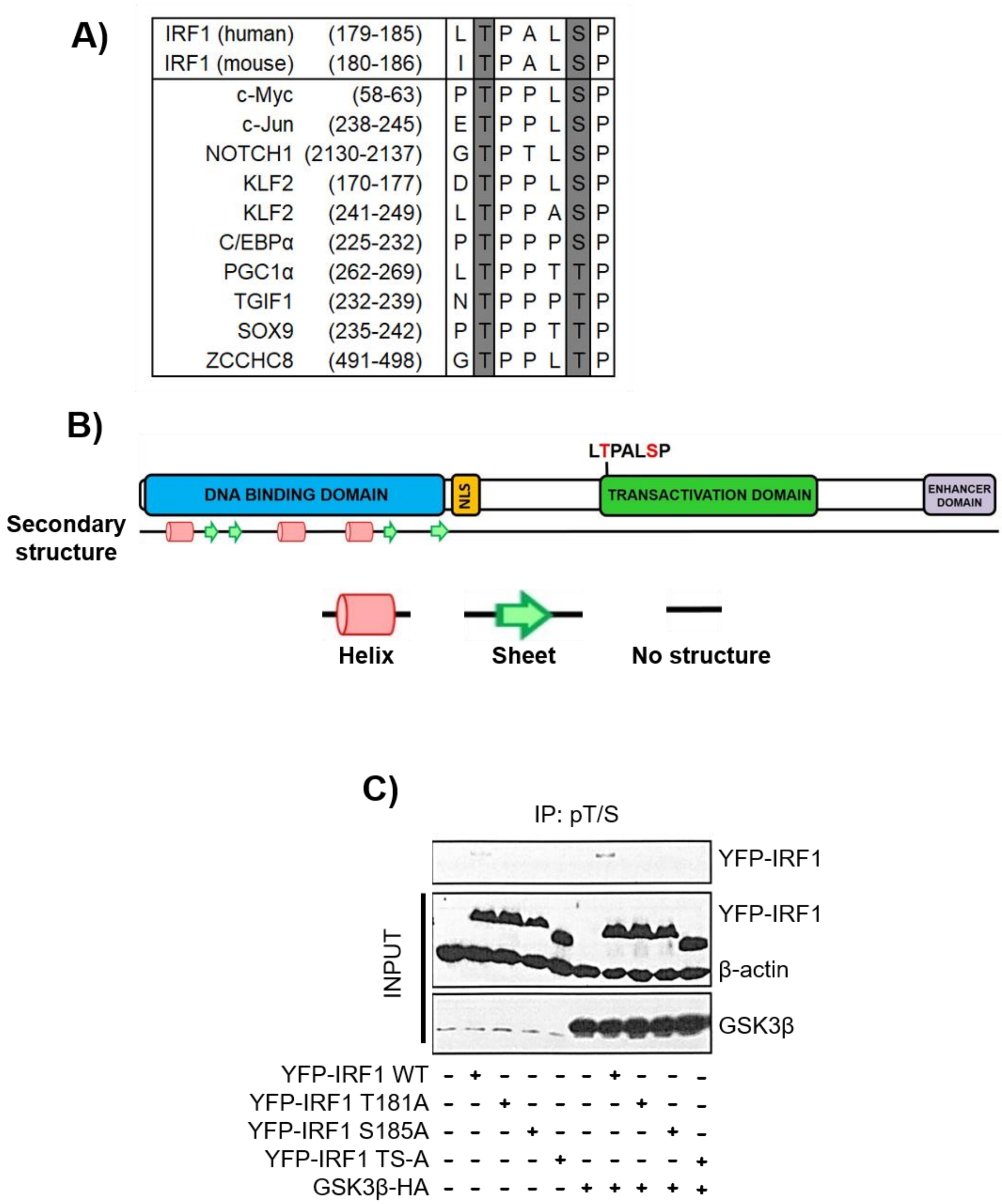

**Figure S2. GSK3β is required for IRF1 transcriptional activity**

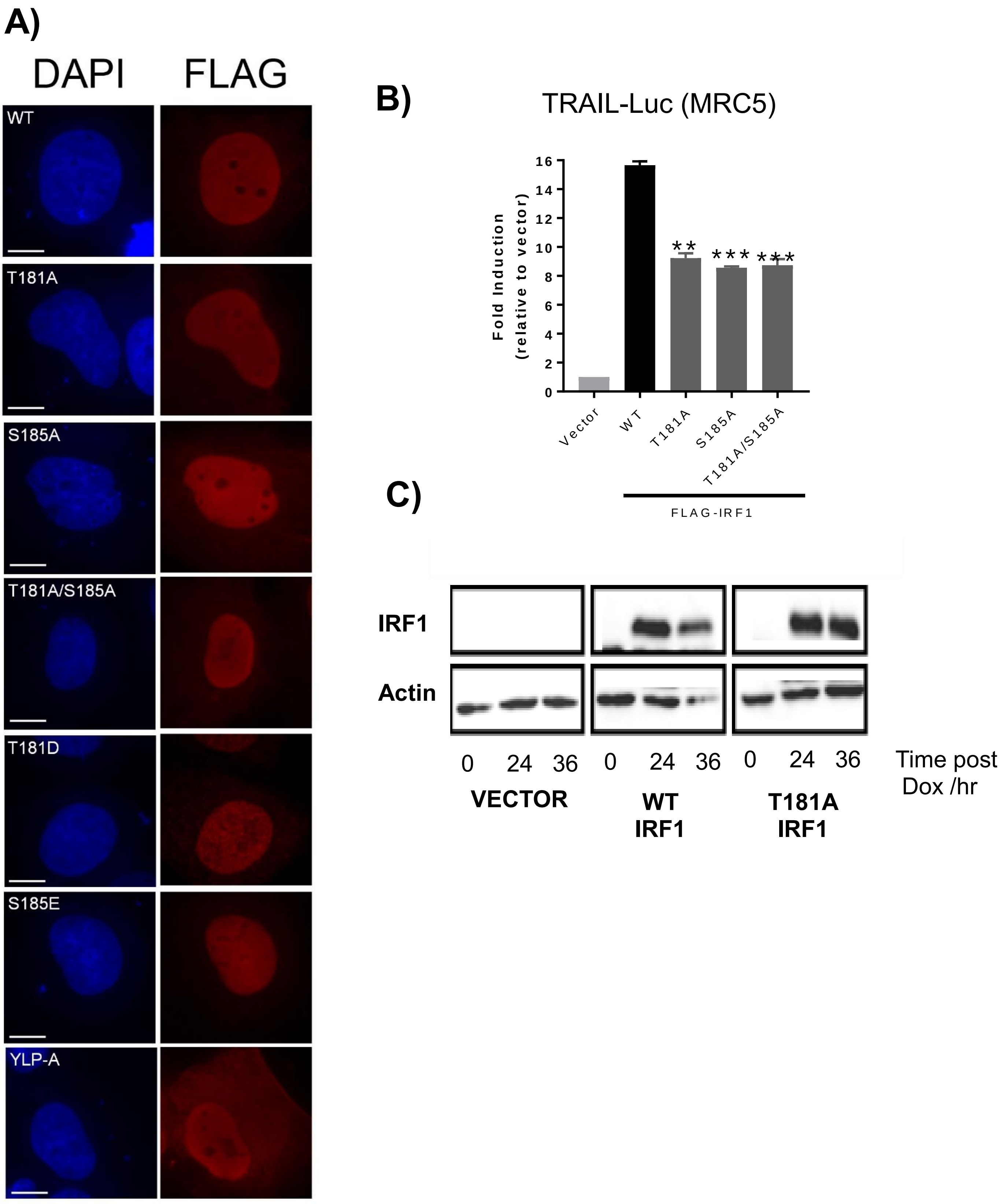

**Figure S3. Stability of IRF1 T181A is not affected by GSK3β over-expression and GSK3β inhibition stabilises IRF1**

A)

| | Half life / minutes | |
|---|---|---|
| | HEK293 | MRC5 |
| WT | 32 | 35 |
| T181A | 60 | 70 |
| S185A | 50 | 55 |
| TS-A | 65 | 75 |
| T181D | 21 | 22 |
| S185E | 22 | 23 |

B)

| | Half life / minutes | |
|---|---|---|
| IRF1 + | HEK293 exogenous mouse IRF1 | MRC5 endogenous human IRF1 |
| vector | 45 | 40 |
| GSK3β WT | 25 | 30 |
| GSK3β K85A | 80 | 70 |

C)

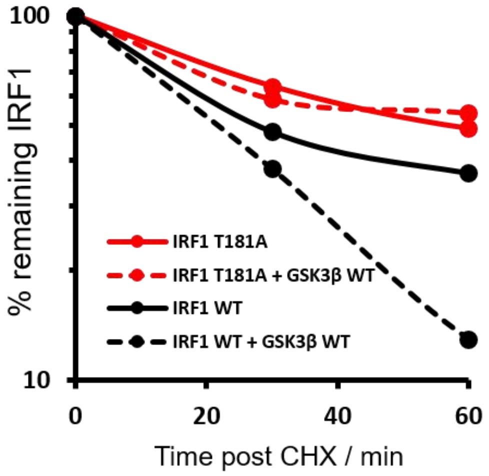


D)

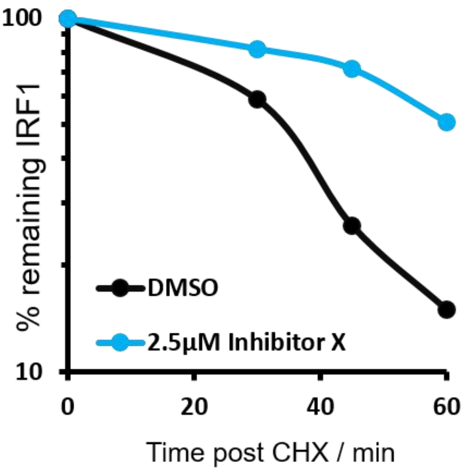

**Figure S4. Fbxw7α interacts with IRF1**

A)

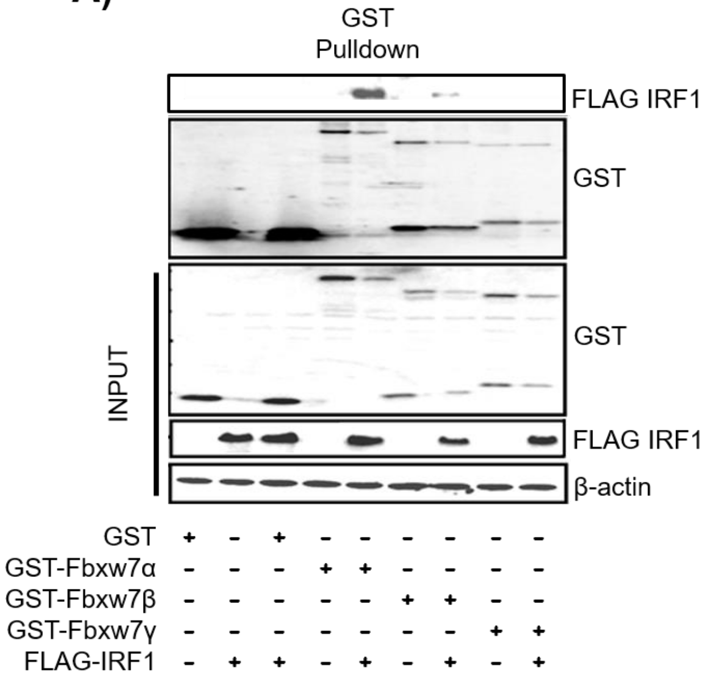


B)

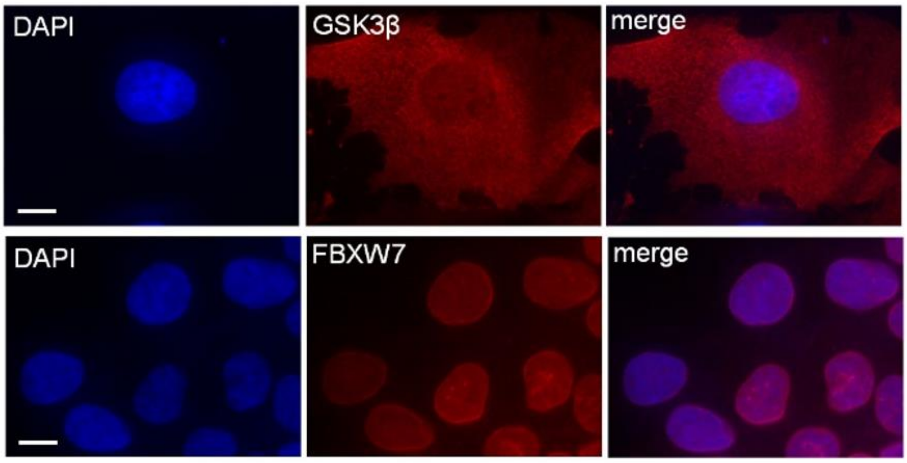

### Figure S5. Fbxw7α regulates IRF1 ubiquitination, half-life and transcriptional activity

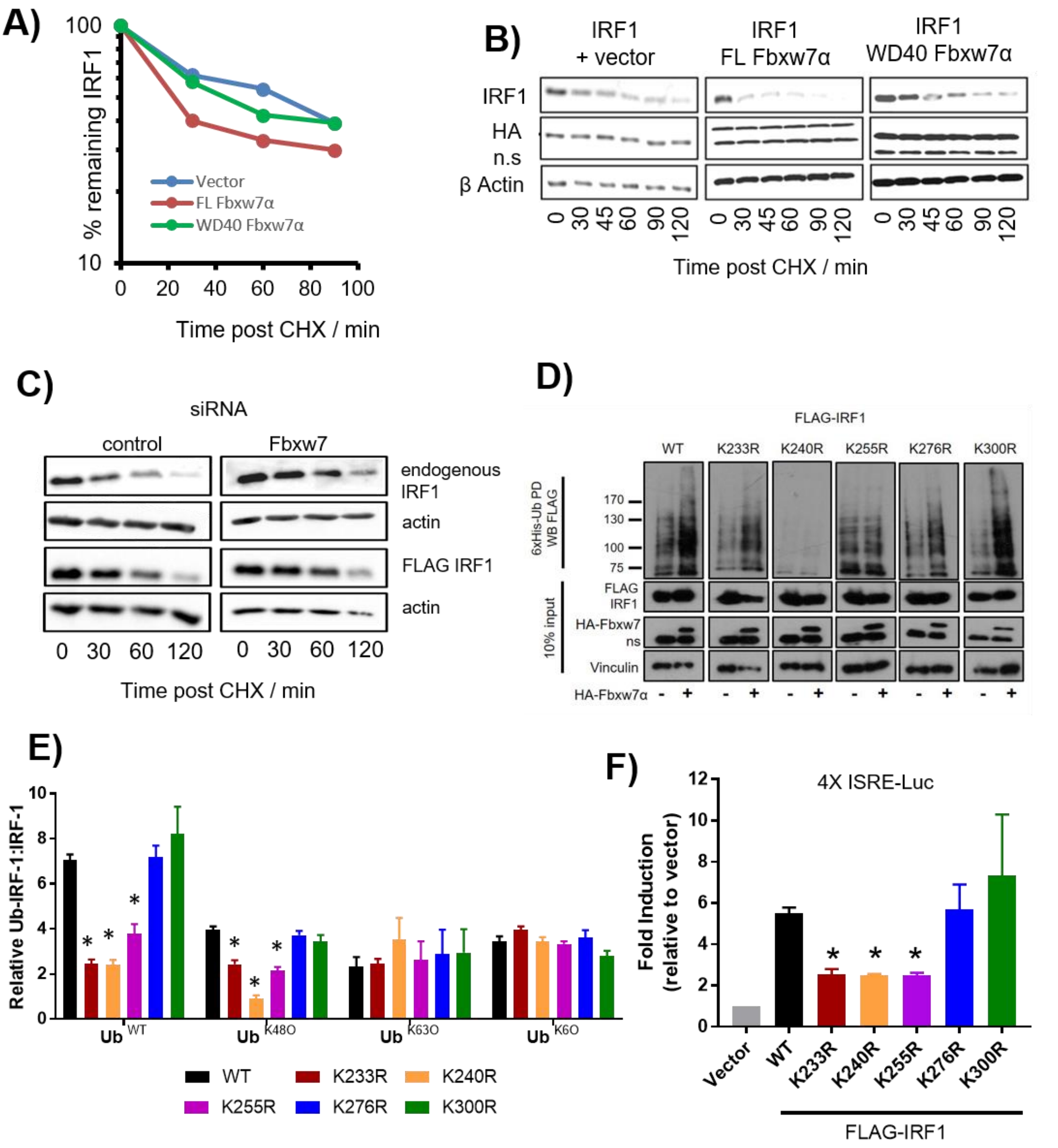

**Figure S6 Thr[181] and Fbxw7 are required for IRF1 anti-proliferative activity in cancer cells.**

**A)**

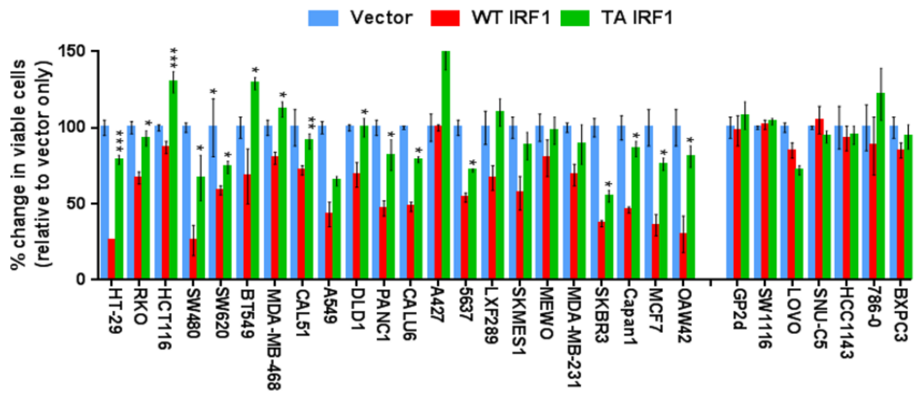

**Supplementary Table 1. Cell lines**

| Cell Line | Tissue | Growth medium | Fbxw7 status* |
|---|---|---|---|
| 5637 | Urinary Bladder | RPMI | WT |
| A427 | Lung | EMEM | WT |
| A549 | Lung | DMEM | WT |
| BT549 | Breast | DMEM | WT |
| CAL51 | Breast | DMEM | WT |
| CAPAN1 | Pancreas | DMEM | WT |
| CALU6 | Lung | EMEM | WT |
| DLD1 | Colon | DMEM | WT |
| HCT116 | Colon | DMEM | WT |
| HT29 | Colon | DMEM | WT |
| LXF289 | Lung | RPMI | WT |
| MCF7 | Breast | DMEM | WT |
| MDA-MB-231 | Breast | DMEM | WT |
| MDA-MB-468 | Breast | DMEM | WT |
| MeWo | Melanoma | EMEM | WT |
| OAW42 | Ovarian | DMEM | WT |
| PANC1 | Pancreas | DMEM | WT |
| RKO | Colon | DMEM | WT |
| SKBR3 | Breast | RPMI | WT |
| SKMES1 | Lung | EMEM | WT |
| SW480 | Colon | DMEM | WT |
| SW620 | Colon | DMEM | WT |
| 786-O | Kidney | RPMI | Homozygous deletion |
| BxPC3 | Pancreas | RPMI | Homozygous deletion |
| GP2d | Colon | DMEM | H580R (WD40 repeats) |
| HCC1143 | Breast | RPMI | Deletion (Mao *et al.* 2010) |
| LoVo | Colon | DMEM | R505C (WD40 repeats) |
| SNU-C5 | Colon | DMEM | S668fs39 |
| SW1116 | Colon | DMEM | H460Y (WD40 repeats) |
| H3396 | Breast | RPMI | WT |
| MRC5 | Lung fibroblast | Alpha MEM | WT |

* Fbxw7 status was determined from the cancer cell line encyclopaedia

**Supplementary Table 2. Antibodies**

| Antibody / Concentration | Supplier | Use |
|---|---|---|
| Murine IRF1 (M20) (1:1000) | Santa Cruz | WB, IP |
| Human IRF1 (C20) (1:1000) | Santa Cruz | WB, IP |
| Phospho-c-Myc ($Thr^{58}/Ser^{62}$) (pT/S) (1:1000) | Santa Cruz | WB, IP |
| GSK3β (27C10) (1:2000)<br>GSK3β ab93926 (1:500) | CST<br>Abcam | WB, IP<br>IF |
| Phospho-Threonine-Proline (9391) (1:1000) | CST | IP |
| GFP (1:2000) | Roche | WB |
| GST (1:2000) | SIGMA | WB |
| FLAG M2 (1:2000) | SIGMA | WB, IP |
| HA 12CA5 (1:2000) | SIGMA | WB, IP |
| β-Actin (A5441) (1:2000) | SIGMA | WB |
| Tubulin (T8203) (1:2000) | SIGMA | WB |
| RNAP II (N20) | Santa Cruz | ChIP |
| pSerine 2 RNAP II (ab5095) | Abcam | ChIP |
| Vinculin (ab129002) (1:2000) | Abcam | WB |
| Ki57 (ab15580) | Abcam | IF |
| Histone H3 (ab10779) (1:2000) | Abcam | WB |
| GAPDH (ab8245) (1:2000) | Abcam | WB |
| Fbxw7 (ab109617) (1:1000) | Abcam | IF |

**Supplementary table 3. Primers**

| Primer | Sequence |
|---|---|
| IRF1 T181A | F: ATGGAAAGGGACATAGCTCCAGCACTGTCACCG<br>R: CGGTGACAGTGCTGGAGCTATGTCCCTTTCCAT |
| IRF1 S185A | F: CATAACTCCAGCACTGACACCGTGTGTCGTCAG<br>R: CTGACGACACACGGTGTCAGTGCTGGAGTTATG |
| IRF1 T181A/S185A | F: GGAAAGGGACATAGCTCCAGCACTGGC<br>R: GCCAGTGCTGGAGCTATGTCCCTTTCC |
| IRF1 T181D | F: GGACTTGGATAGGAAAGGGACATAGATCCAGCACTGTCA<br>R: TGACAGTGCTGGATCTATGTCCCTTTCCATATCCAAGTCC |
| IRF1 S185E | F: AGGGACATAACTCCAGCACTGGAGCCGTGTGTCGTCAGCAGCAGT<br>R: TCCCTGTATTGAGGTCGTGACCTCGGCACACAGCAGTCGTCGTCA |
| IRF1 YLP-A | F: GCGGGTGGCCCGGATGGCGGCACCCCTCACCAGG<br>R: CCTGGTGAGGGGTGCCGCCATCCGGGCCACCCGC |
| IRF1 K233R | F: GGATGAGGAAGGGAGGATAGCCGAAGACC<br>R: GGTCTTCGGCTATCCTCCCTTCCTCATCC |
| IRF1 K240R | F: GATAGCCGAAGACCTTATGAAGGCTCTTTGAACAGTCTGAG<br>R: CTCAGACTGTTCAAAGAGCCTCATAAGGTCTTCGGCTATC |
| IRF1 K255R | F: GACACACATCGATGGCAGGGGATACTTGCTCAATG<br>R: CATTGAGCAAGTATCCCCTGCCATCGATGTGTGTC |
| IRF1 K276R | F: GGAGACTTCAGCTGCAGAGAGGAACCAGAGATTG<br>R: CAATCTCTGGTTCCTCTCTGCAGCTGAAGTCTCC |
| IRF1 K300R | F: CATGTCTTCACGGAGATGAGGAATATGGACTCCATCATG<br>R: CATGATGGAGTCCATATTCCTCATCTCCGTGAAGACATG |
| IRF1 (pEYFP) | F: ATAATAAGATCTATGCCAATCACTCGAATG<br>R: ATAATATCTAGACTATGGACAAGGAAT |
| IRF1 (FLAG) | F: ATAATAAAGCTTATGCCAATCACTCGAATG<br>R: ATAATATCTAGACTATGGACAAGGAAT |
| HA-Fbxw7α FL | F: ATAATAGAATTCATGAATCAGGAACTGCTCTCTGTG<br>R: TATTATTCTAGATCACTTCATGTCCACATCAAAGTC |
| HA-Fbxw7α FL ΔWD40 | F: ATAATAGAACCTAAGGTGCTGAAAGGACATGAT<br>R: TATTATTCTAGATCAAGATTTGAGTTCTCCTCGCCT |
| RT-QPCR primers | Clarke *et al.* 2004 |
| ChIP primers | Clarke *et al.* 2004 |
